# Metabolic growth coupling strategies for *in vivo* enzyme selection systems

**DOI:** 10.1101/2024.09.16.613188

**Authors:** Tobias B. Alter, Pascal A. Pieters, Colton J. Lloyd, Adam M. Feist, Emre Özdemir, Bernhard O. Palsson, Daniel C. Zielinski

## Abstract

Whole-cell biocatalysis facilitates the production of a wide range of industrially and pharmaceutically relevant molecules from sustainable feedstocks such as plastic wastes, carbon dioxide, lignocellulose, or plant-based sugar sources. The identification and use of efficient enzymes in the applied biocatalyst is key to establishing economically feasible production processes. The generation and selection of favorable enzyme variants in adaptive laboratory evolution experiments using growth as a selection criterion is facilitated by tightly coupling enzyme catalytic activity to microbial metabolic activity. Here, we present a computational workflow to design strains that have a severe, growth-limiting metabolic chokepoint through a shared class of enzymes. The resulting chassis cell, termed enzyme selection system (ESS), is a platform for growth-coupling any enzyme from the respective enzyme class, thus offering cross-pathway application for enzyme engineering purposes. By applying the constraint-based modeling workflow, a publicly accessible database of 25,505 potential and experimentally tractable ESS designs was built for *Escherichia coli* and a broad range of production pathways with biotechnological relevance. Model-based analysis of the generated design database reveals a general design principle that the target enzyme activity is linked to overall microbial metabolic activity, not just the synthesis of one biomass precursor. Furthermore, the use of currently available genome-reduced strains or one preeminent carbon source does not significantly enable or improve growth-coupling of target enzymes. Most importantly, observed trade-offs between the predicted viability of ESSs, the design-inflicted metabolic perturbations, and the coupling strength suggest that a suboptimal coupling has benefits regarding the experimental implementation of ESSs and growth-coupling in general. Overall, the computed design database, which is accessible through https://biosustain.github.io/ESS-Designs/, and its analysis lay the foundation for generating valuable *in vivo* ESSs for a range of biotechnological applications.

Graphical abstract

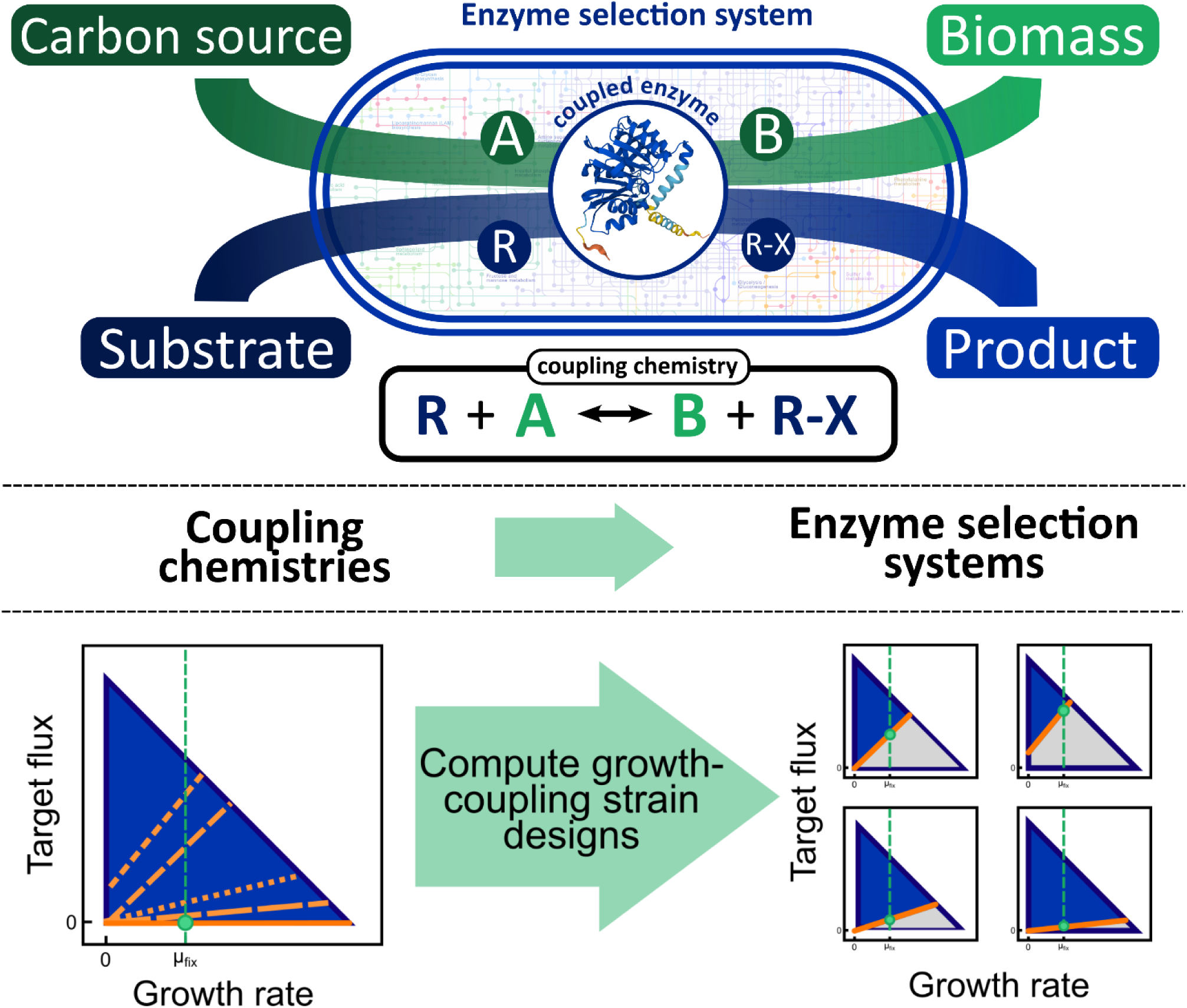

**Highlights:**

- Enzyme selection system designs link enzyme activities to global metabolic processes.
- Enzymes can be categorized based on growth-couplable reaction stoichiometries.
- A database of strain designs for building enzyme selection systems was created.
- Enzyme selection systems enable optimization of a range product synthesis pathways.
- Application recommendations: weakly coupled enzyme activities can have practical advantages.

## 1. Introduction

Microbial production of value-added molecules in integrated bioprocesses is widely recognized as a major driver towards a sustainable, circular bioeconomy. The microorganisms used in such bioprocesses typically require significant engineering to achieve economically feasible yields and production rates under fermentation conditions. Genetic engineering efforts usually involve the expression of heterologous enzymes enabling a microbe to synthesize a target molecule from its native metabolites. The performance of heterologous enzymes, i.e., the maximum achievable rate with which substrates are converted to products, is a key factor determining the performance of the overall production pathway.

Neither heterologous nor native enzymes are typically adapted for maximum production efficiency in an engineered microbial host (Bar-Even et al., 2011). Traditional enzyme engineering techniques, such as random mutagenesis or sequence-based semi-rational strategies (Cheng et al., 2015), can be applied to improve relevant enzyme properties, but require significant experimental investment. Generation and screening of enzyme mutant libraries need to be conducted at high throughput, which necessitates non-standard laboratory equipment and an assay suitable for rapid and reliable testing of the target property (Shivange et al., 2009). Effective screening of large enzyme libraries is becoming more accessible due to the worldwide advance of biofoundries (Hillson et al., 2019). However, establishing proper screening assays remains a non-trivial and laborious task. Moreover, assays are often conducted *in vitro*, thus may not resemble the native biophysical conditions enzymes are subjected to within a cell, potentially leading to unfavorable discrepancies in the performance of a selected enzyme variant between the *in vitro* screening and the *in vivo* application conditions.

*In vivo* enzyme screening and selection systems based on a strict metabolic coupling between a target enzyme’s activity and growth, here referred to as enzyme selection systems or ESSs, have several advantages. ESSs allow for the use of growth as a simple selection criterion for target enzyme activity. This feature is exploited for enzyme engineering purposes in Adaptive Laboratory Evolution (ALE) experiments during which a microbial culture is repeatedly propagated into fresh medium and naturally selected for favorable mutations leading to higher growth rates (Sandberg et al., 2019). The necessity for screening assays or high throughput instrumentation is omitted because cells harboring favorable mutations will outgrow other with detrimental or neutral mutations. ALE is thereby not restricted to adaptations in the target enzyme’s primary sequence but can simultaneously select for, as an example, an improved cofactor or precursor supply in the microbial host. As a proof-of-principle for ESSs, an *Escherichia coli* strain has recently been designed to couple methyltransferases to growth and subsequent ALE experiments have led to improved enzymes variants (Luo et al., 2019).

Due to the vast complexity and redundancy of metabolic networks, a generalized approach for growth-coupling any enzyme requires computer-aided techniques to derive appropriate genetic designs. Various computational methods that use constraint-based metabolic models as dense biochemical knowledgebases have been built to serve this purpose (Alter and Ebert, 2019; Burgard et al., 2003; Feist et al., 2010; Jensen et al., 2019; Pharkya et al., 2004; von Kamp and Klamt, 2014). While most model-based studies have focused on computing genetic designs for coupling growth to the production of a target molecule or the activity of endogeneous reactions, growth-coupling of specific enzymes has not been extensively considered.

In this study, we developed chassis ESS designs in *E. coli* for a range of enzyme classes. Each single chassis ESS is applicable to all enzymes within the associated enzyme class, thus representing a “plug-and-play” *in vivo* engineering and optimization tool for multiple enzymes with a single strain design. For this purpose, we defined an appropriate enzyme class as a group of enzymes sharing a common substrate-product pair, which we term a coupling chemistry (CC). Linking a CC to growth produces an ESS for all enzymes sharing that CC due to their common substrate-product pair. We identified an exhaustive list of CCs for *E. coli* and selected for candidates which allowed for an integration into the genome-scale constraint-based metabolic model iML1515 (Monk et al., 2017). Furthermore, a computational framework was established and applied for identifying diverse sets of growth-coupled strain designs for each modeled CC considering gene knockouts, heterologous reaction insertions, carbon source, and metabolite cofeeds as engineering variables. We show that ESS strain designs exist for 33 CCs, and that a notable number of CCs hold the potential for ESSs with a strong selection pressure. The enzymes in the subset of CCs with feasible growth-coupled strain designs span various metabolic pathways. This finding supports the potential universality of *in vivo* ESSs as a tool for optimizing enzymes towards efficient, high-flux target molecule synthesis routes. Generally, CCs involved in the transfer of a high-molecular group, such as a CoA cofactor, show a decreased success rate in finding strong coupling strain designs compared to those transferring low-molecular groups. Moreover, the target enzyme activity in most identified designs is linked to the overall microbial metabolic activity, not just the synthesis of one biomass precursor, thus implies to yield tighter growth-coupling. All identified strain designs, performance metrics such as predicted maximum growth rates and coupling strengths, and value-added target molecules associated with the designs or CCs are made available online via https://biosustain.github.io/ESS-Designs/.

## 2. Methods

### 2.1. Software, hardware, and test statistics

All workflow and analysis code was implemented and tested in Python 3.8 using recent versions of juypterlab ≥ 2.2.4 (Jupyter Development Team, 2020), numpy ≥ 1.19.1 (Harris et al., 2020), pandas ≥ 1.4.0 (The pandas development team, 2020), scipy ≥ 1.5.0 (Virtanen et al., 2020), cobra ≥ 0.20.0 (Ebrahim et al., 2013), cameo ≥ 0.13.6 (Cardoso et al., 2018), optlang ≥ 1.5.1 (Jensen et al., 2017), gurobipy ≥ 9.1.2 (Gurobi Optimization, 2023), equilibrator-api ≥ 0.3.2 (Beber et al., 2021), and related packages. All data plots were made with matplotlib ≥ 3.3.1 (Hunter, 2007) and matplotlib-venn ≥ 0.11.6 (Tretyakov, 2022).

All Kolmogorov-Smirnov tests were performed using the ks_2samp function from scipy’s statistical functions module. Pearson correlation statistics were retrieved from scipy’s linear least-squares regression function linregress.

Simulations and analyses as part of the workflow described in the following section were conducted on the high-performance computing cluster of the Technical University of Denmark (DTU) using a maximum configuration of 48 GB of RAM and 24 Intel Xeon 2660v3 processors (2.60 GHz). A Windows 10 machine with 32 GB of RAM and an AMD Ryzen 5900x processor (12 cores at 3.7 GHz) was used for data processing.

### 2.2. A metabolic model-based workflow for computing growth-coupling strain designs

A Python-based workflow was implemented for computing and evaluating growth-coupling strain designs by exploiting the rich metabolic information content of constraint-based metabolic models. COBRA toolbox functionalities were integrated to enable intuitive and community-friendly model management. The workflow can be divided into three principal steps: (i) set up of the metabolic model and optimization algorithm parameters, (ii) search for growth-coupling strain design solutions using the gcOpt algorithm (Alter and Ebert, 2019), and (iii) evaluation and summarization of design results. The Python code for running the workflow, here termed the Growth-Coupling Suite, is freely available online on https://github.com/biosustain/Growth-coupling-suite. Examples are provided in Jupyter notebooks to guide the user through typical workflows. While this study focuses on designing and evaluating growth-coupled strains using coupling chemistries (CCs) in the *E. coli* model, this workflow is applicable to any target reaction or metabolic model.

#### 2.2.1. Metabolic model and optimization algorithm set up

Setting up the metabolic model and solver parameters is a crucial first step in the general workflow. All specific parameters used to compute growth-coupling strain designs for ESSs in this study are given in the following section. Environmental conditions, i.e., medium components such as the carbon source, or a strain-specific genotype in terms of gene knockouts and heterologous reactions are applied to the metabolic model by properly setting relevant reaction bounds. Prior knowledge from single-gene knockout or transposon insertion sequencing studies can be considered for constraining the design solutions space by defining essential, thus non-deletable metabolic genes or reactions. To allow for the consideration of new metabolic network edges as design variables, a diverse set of heterologous reactions is derived from 28 genome-scale metabolic models for different species taken from the BIGG model database (Schellenberger et al., 2010) including the *Escherichia*, *Saccharomyces*, *Mycobacterium*, *Homo*, and *Bacillus* genera among others (Table S1). A detailed description of generating databases of heterologous reactions is given in section 2.4. To allow for the optimization of the medium composition for generating growth-coupling designs, exchange reactions of potential carbon or cofeed sources are declared as design variables. Finally, optional solver- and optimization problem-specific parameters such as maximum computation times, maximum number of total and specific interventions, or number of CPU processes to be used are set.

Before solving the gcOpt problem using the prepared model, the number of heterologous reaction insertions and gene deletion target variables are minimized without eliminating possible solutions. A minimal yet meaningful set of design variables ensures time-efficient solving of the optimization problem. Flux variability analysis (FVA) is applied to detect blocked reactions which cannot carry flux under any admissible environmental condition (Mahadevan and Schilling, 2003). During FVA, the uptake of all metabolites defined as potential carbon sources is simultaneously enabled by assigning negative values to the lower bounds of their corresponding exchange reactions. A reaction is blocked, if its maximum and minimum flux is zero. Blocked heterologous and native reactions are erased from the model, thus removed as design variables. If a reaction is enforced, i.e., its flux rate takes a non-zero value under any given condition, its associated genes are disregarded as knockout candidates as well. Note that all isozymes catalyzing a blocked or enforced reaction are excluded from the design space, since information about the predominance in catalytic activity among isozymes is scarce and deciding which enzyme deletion may not affect a rection’s activity is not easily made *a priori*.

#### 2.2.2. Computing growth-coupling strain designs

At the core of the workflow, a gcOpt algorithm (Alter and Ebert, 2019) implementation searches for strain designs which maximize a minimal guaranteed flux rate of a target reaction at a fixed growth rate. For a detailed description of gcOpt and the underlying optimization problem formulation refer to Alter and Ebert (2019). In contrast to the original gcOpt implementation, heterologous reaction insertions and medium conditions can additionally be considered as design variables besides reaction knockouts. The corresponding bilevel Mixed-Integer Linear Programming (MILP) problem is expressed mathematically as follows:

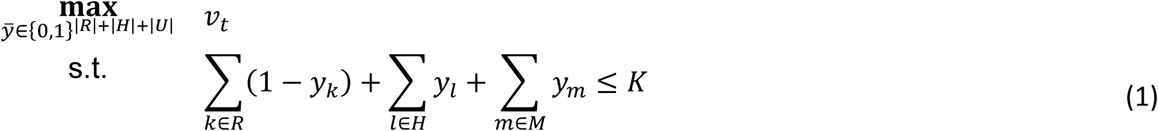

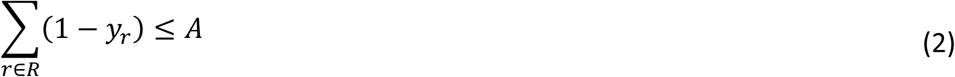

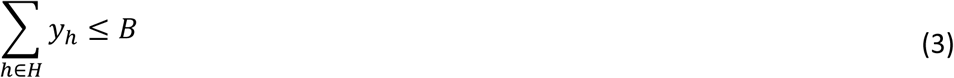

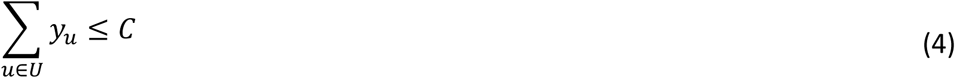

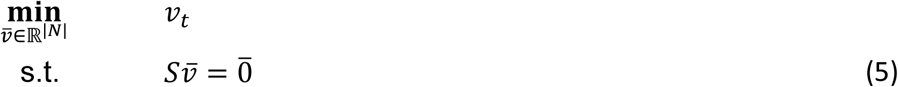

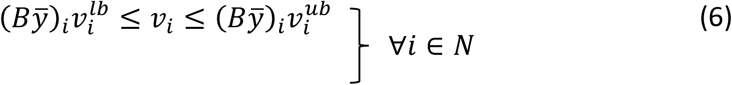

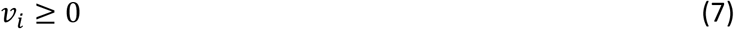

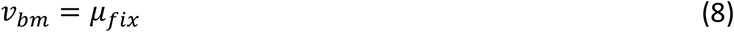

Here, *y̅* is a Boolean vector indicating if a target reaction is active (1) or inactive (0). The decision variables in *y̅* are divided into separate sets *R*, *H*, and *U* representing knockouts, heterologous insertions, and substrate uptake reactions, respectively. The total number of interventions is restricted to *K* (Eq. 1). The number of knockouts, heterologous reaction insertions, and substrate uptake reactions are separately limited to *A*, *B*, and *C*, respectively (Eq. 2-4). Inactivation of a target reaction *i* is enforced by setting its lower and upper bound to zero, if *y*_*i*_ = 0 (Eq. 6). Otherwise, the native lower and upper bounds 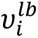 and 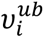 are active. Note that the MILP formulation requires an irreversible metabolic network where reaction fluxes are strictly positive (Eq. 7). Thus, reversible reactions are split into forward and backward reactions resulting in a metabolic model with *N* irreversible reactions. Consequently, a matrix *B* is introduced to map decision variables of each reversible reaction to their irreversible counterparts (Eq. 6). Fluxes of each irreversible reaction are contained in *v̅*. The metabolic network is represented by the stoichiometric matrix *S* ∈ ℝ^|*M*|+|*N*|^ containing the stoichiometric coefficients of all metabolites *j* ∈ *M* participating in reactions *i* ∈ *N*. Steady state mass balances of all metabolites within the metabolic network are ensured by Eq. 5. The growth rate represented by the flux through the biomass formation reaction *v*_*bm*_ is fixed to *μ*_*fix*_ by Eq. 8. Note that the fix growth rate value *μ*_*fix*_ acts like a lower bound on the maximum growth rate *v*_*bm*,*max*_. Thus, predicted maximum growth rates higher than *μ*_*fix*_ are still possible for gcOpt design solutions, since flux solutions with *v*_*bm*_ = *μ*_*fix*_ are always contained in the convex solution space of a (perturbed) metabolic network for which *v*_*bm*,*max*_ ≥ *μ*_*fix*_ holds.

The gcOpt MILP problem is solved by the Gurobi Optimizer suite (version 9.0.0 or higher) (Gurobi Optimization, 2023) providing suboptimal solutions if optimality is not reached in practical solving times. Since feasible solutions guarantee a non-zero target reaction flux for a range of growth rates, the corresponding strain designs enforce growth-coupling on the target reaction. Moreover, model inherent Gene-Protein-Reaction (GPR) relations can be used at solver run-time through the Gurobi callback class to validate the feasibility of reaction knockouts on the gene level. That is, if a set of reaction knockouts in a strain design solution cannot be realized with gene knockouts, the solution is made infeasible by adding appropriate lazy constraints to the MILP problem before optimization is continued.

#### 2.2.3. Evaluation of computed design results

After termination of the gcOpt algorithm, either upon reaching the optimal design solution or the time limit, each identified strain design is evaluated using the original linear program model. Primarily, maximum growth rates under optimized medium conditions and the growth-coupling strength (GCS) are computed by maximizing the biomass formation flux. The GCS of growth-coupling strain designs is computed as described in section 2.5. These values, along with other metadata such as genetic interventions or the carbon source used, are stored are stored in an Excel file for each strain design output by gcOpt. Relevant designs can be selected from the summary file based on design complexity (e.g., the number of genetic interventions), predicted coupling strength, or expert knowledge.

### 2.3. Computing enzyme selection system strain designs

An enzyme selection system (ESS) comprises a CC coupled to microbial growth. To investigate the scope of application for ESSs in *E. coli*, growth-coupling strain designs were computed for various selected CCs using the workflow described in the previous section. For each CC, the workflow was applied multiple times with seven models representing different genetic or environmental conditions (Table 1). The genome-scale metabolic model iML1515 for *E. coli* K12 MG1655 (Monk et al., 2017) taken from the BIGG model database (Schellenberger et al., 2010) was used as base model.

**Table 1.** Model variants or parameter adaptions applied to compute growth-coupling strain designs for each of the 44 selected coupling chemistries. All models base on the “Standard” parameter set and were adapted as described. The purpose of applying a model, i.e., the research question that is to be answered, is additionally given.

| Model ID | Model specifics | Purpose |
| --- | --- | --- |
| Standard | iML1515 under aerobic conditions with model and optimization parameters as described in the Methods section | Use the full metabolic capability of *E. coli* and all common design variables. |
| No cofeed | Disabled selection of a metabolite cofeed in addition to a carbon source | Does the use of an external metabolite cofeed promote the establishment of growth-coupling? |
| N-coupling | Inactivated ammonia exchange to remove the primary nitrogen source | Does an enforced nitrogen supply via a cofeed metabolite and the target reaction lead to better growth-coupling strategies? |
| Anaerobic | Inactivated oxygen exchange to model anaerobic conditions | Is growth-coupling favored under anaerobic conditions? |
| MS56 | Deleted genes from iML1515 to match the genome-reduced strain MS56 | How does genome reduction influence the design of growth-coupling strategies? |
| DGF-298 | Deleted genes from iML1515 to match the genome-reduced strain DGF-298 |  |
| Essential genes | Allow for the deletion of known essential genes | Does the essentiality of a few genes hinder the realization of growth-coupling strain designs? |

#### 2.3.1. Applying environmental and genetic conditions to the metabolic model

In a first step, iML1515 was adapted to match the characteristics of the respective models or conditions described in Table 1. For the N-coupling and Anaerobic case, the lower bound of the NH_4_ and O_2_ exchange reactions were set to 0 to disable the uptake of ammonia and oxygen, respectively. In all other cases the oxygen uptake rate was limited to 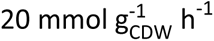 (Varma et al., 1993) and the ammonia uptake was effectively unbounded 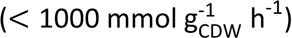. The uptake of any carbon source was initially disabled, since the choice for a carbon source was a design variable in any case, and thus subject of the gcOpt optimization problem. The solver was forced to activate the uptake of one of 48 predefined carbon sources (Supplementary Data) in a design solution. Except for the No-Cofeed case, an optional uptake of one other metabolite (cofeed) was additionally allowed. Any exchange reaction for a metabolite with maximum 11 carbon atoms were considered as a cofeed reaction candidates. 33 metabolites with higher carbon contents were explicitly added as cofeed candidates (Supplementary Data). Throughout this study, the maximum uptake rate for each carbon or cofeed source was constrained to 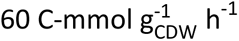 and 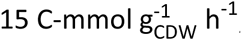, respectively. For the MS56 and DGF-298 case, metabolic genes were deleted from iML1515 to match the reduced genomes of the MS56 (Park et al., 2014) and DGF-298 (Hirokawa et al., 2013) strains, respectively. The lists of deleted genes can be found in the Supplementary Data. Reaction deletions corresponding to the sets of deleted genes were derived from the gene-protein-reaction (GPR) relations given in iML1515.

#### 2.3.2. Integration of the coupling chemistry into the metabolic model

Secondly, generic stoichiometric reactions for the respective CC (Table 2) was added to the metabolic model. This reaction was set as the target or objective in the gcOpt problem. Each reaction consists of a specific substrate-product pair native to the model, as well as a generic substrate and product compound representing the various reactants annotated for enzymes corresponding to the CC. The elemental compositions of the generic compounds were adapted to close the mass balance of the reaction. The reaction was set as irreversible into the forward direction with the positive flux rate being effectively unbounded 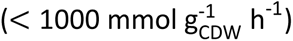. Exchange reactions for the generic compounds were added to the model to allow for their free balancing. Transport costs for the generic compounds over the membranes were neglected, since these cannot be generalized across the broad spectrum of known compounds and transport mechanisms. An energy (ATP) or proton gradient demand of metabolite transport across membranes can affect or even enable growth coupling according to a theoretical study using an *E. coli* core metabolic model (Alter and Ebert, 2019), thus requires consideration when evaluating computed strain designs for enzyme- and reaction-specific case examples.

**Table 2.**
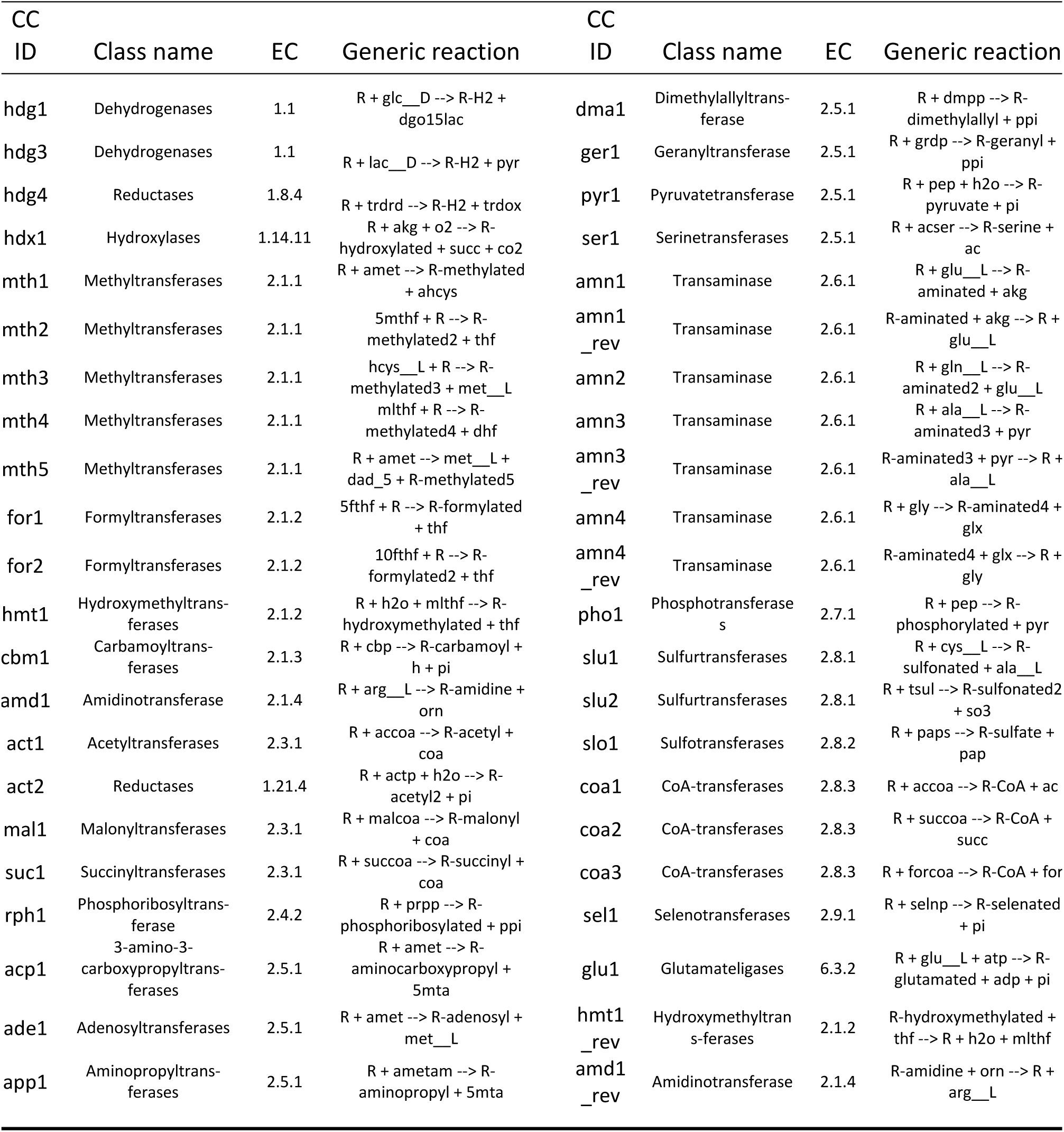
44 coupling chemistries (CCs) selected for growth-coupling simulations in the *E. coli* metabolic model iML1515 to derive enzyme selection system designs. R represents a generic compound in the reaction stoichiometries and is a placeholder for an enzyme-specific reactant. All other compound abbreviations follow the standard nomenclature from the BIGG database. Note that a given EC annotation represents the majority and not necessarily all enzymes within the corresponding CC.

#### 2.3.3. Reduction of the design variable solution space and set up of optimization parameters

Besides the general mathematical techniques to reduce the solution space of the gcOpt problem (cf. section 2.2.1), biological logic and data were used to further reduce the number of design variables. Genes observed to be essential for *E. coli* were removed from the solution space as design variables. The list of essential genes is given in the Supplementary Data and was derived from EcoCyc (Keseler et al., 2021) and the Profiling of *E. coli* Chromosome (PEC) database (Yamazaki et al., 2008) where gene essentiality data was collected from various sources (Baba et al., 2006; Feist et al., 2007; Joyce et al., 2006; Patrick et al., 2007). Although the gene essentiality data used are conditional, they provide a good indication of gene essentiality in minimal medium under standard conditions for each substrate, taking into account to the redundancy of the metabolic network. Furthermore, a selection of reactions associated with the electron transport chain or specific metabolic subsystems were additionally excluded as knockout targets. The selection includes NADH dehydrogenases (NADH16pp, NADH17pp, NADH18pp), NAD transhydrogenase (NADTRHD), ATP synthase (ATPS4rpp), cytochrome oxidases (CYTBO3_4pp, CYTBDpp, CYTBD2pp), and reactions assigned to the subsystems Cell Envelope Biosynthesis, Inorganic Ion Transport and Metabolism, Lipopolysaccharide Biosynthesis / Recycling, Murein Biosynthesis, Murein Recycling, tRNA Charging, as well as all transport reactions.

To derive diverse sets of growth-coupling strain designs for each CC-model combination, multiple optimization parameter sets were applied independently for any given gcOpt problem. The total number of interventions including heterologous reaction insertions, gene knockouts, and the addition of a metabolite cofeed was set to 5, 8, and 11 in separate gcOpt optimization runs. The decision for the exact distribution of intervention types (primarily insertions and knockouts) in a design solution was left to the solver, but the number of cofed metabolites was always restricted to one. Furthermore, two fixed growth rates (0.1 and 0.3 ℎ^−1^) were applied adding up to six independent gcOpt optimization problems for each CC-model combination which results were combined for further analyses after termination (cf. previous section). Any optimization was run for maximum 10 h and used up to seven processes in parallel.

### 2.4. Generation of heterologous reaction databases

A heterologous reaction database is generated for a target metabolic model by merging reactions from a user-defined set of metabolic reconstructions. Only heterologous reactions that do not overlap in functionality or stoichiometry with any native reaction in the target metabolic model are included in the database. Each reaction undergoes a consistency check before being considered for a heterologous reaction database.

Firstly, the identifier of each participating metabolite was, if needed, adapted to the BIGG (Schellenberger et al., 2010) namespace by consulting the MetaNetX (Moretti et al., 2016) and KEGG (Kanehisa et al., 2017) databases, followed by an update of its elemental composition. If the heterologous reaction was not mass or charge balanced, it was dismissed.

Secondly, the thermodynamics, i.e., the directionality of a reaction, was evaluated. The Equilibrator database (Beber et al., 2021) was consulted to obtain thermodynamic data for defining a reaction as reversible or irreversible in the backward or forward direction. Standard Gibbs energies of reaction Δ_r_*G*^0^were retrieved via the Equilibrator API (Python package equilibrator-api) and transformed with metabolite concentration information to estimate feasible ranges of a reaction’s Gibbs free energy Δ_r_G. Intracellular metabolite concentrations were assumed to range between 0.01 mM and 20 mM for that purpose (Feist et al., 2007). Since concentrations of the cofactors ADP, ATP, AMP, NAD^+^, NADH, NADP^+^, and NADPH are highly regulated in *E. coli*, thus approximately constant across conditions, these were fixed to physiologically relevant values (Table S2) in this study. If the range of Δ_r_G was strictly below or above zero, the heterologous reaction was assigned as irreversible in the forward or backward direction, respectively. If Δ_r_G can take both positive and negative values, the reaction was reversible. Eventually, a heterologous reaction was only accepted as a design variable if the computed reaction direction matched the reaction’s reversibility in the native metabolic model to fortify its thermodynamic consistency.

Identifiers for genes in the host organism corresponding to a heterologous reaction are indicated in the final database. However, the promiscuity of enzymes catalyzing a heterologous reaction is not taken into account. The selection of a specific enzyme for a suggested heterologous reaction within a growth-coupling strategy is ultimately left to the user, as many other factors such as activity, optimal temperature, or inhibiting factors may need to be considered. Therefore, a proper post-processing pipeline is required to derive suitable genetic elements for a growth-coupling strategy. However, this is beyond the scope of this study.

For this work, the heterologous reaction database was built from 28 metabolic models taken from the BIGG database (Table S1). The final heterologous reaction databases for the wildtype *E. coli* model iML1515, the genome-reduced MS56, and DGF-298 models comprise 1298, 1448, and 1523 reactions, respectively. The reactions and corresponding enzymes contained therein are a representative subset of all known enzymes according to their EC number distribution (Fig. S1).

### 2.5. Computing the coupling strength of growth-coupling strain designs

The growth-coupling strength (GCS) is a measure for the interdependency between growth rate and the target reaction flux, here the flux through the CC reaction, and was introduced by Alter and Ebert (2019). Technically, a GCS value depicts the ratio between the feasible flux space of the wildtype and the flux space made inaccessible by the strain design in 2D flux space projections span by the biomass formation and target reaction (Fig. S2). Thus, the GCS quantifies the number of unwanted flux states eliminated by a growth-coupling strain design, i.e., flux states with low or zero target reaction flux at any feasible growth rate. In contrast to the original GCS formulation of Alter and Ebert (2019), the ratio between the minimally guaranteed product yield of the target reaction at maximal growth and the theoretical maximum yield was not considered as an additional factor. The inclusion of a notion of maximum productivity is valid when the synthesis and secretion of a value-added molecule is targeted for growth-coupling but is superfluous for enzyme selection systems where the focus is on ensuring coupling for a range of growth rates. Moreover, omitting the maximum productivity factor reduces the computational demand for calculating GCS values. The GCS ranges between 0 (no growth-coupling) and 1, where increasing values indicate tighter growth-coupling. However, since designs with a GCS of close to 1 would effectively block growth, a relevant maximum GCS for any target reaction is denoted by a value of approximately 0.6.

### 2.6. The ATP synthesis capability of metabolic reactions

The ATP synthesis capability (ATPsc) describes the influence of a reaction flux on the overall production of energy equivalents in the form of ATP in a constraint-based metabolic model. ATPsc for a reaction *i* is defined as the derivative *d*υ_ATP_⁄*d*υ_i_. Positive ATPsc values for *i* denote that the maximum ATP synthesis rate *v*_*ATP*_increases with higher flux rates of *i* and *vice versa*. If the ATPsc is negative, the maximum ATP synthesis decreases with elevated activity of *i*. As shown by Alter and Ebert (2019), ATPsc values computed for exchange reactions of central metabolites in *E. coli* metabolic models reproduced the expected energy hierarchy of metabolites. Klicken oder tippen Sie hier, um Text einzugeben.Acetate shows the highest, positive ATPsc values after CO_2_ which is in line with observed fermentation byproduct secretion as a consequence of limitations in respiratory ATP production (Clark, 1989).

In this study, the ATPsc for a reaction *i* is computed from two linear programs maximizing the unbounded ATP maintenance requirement reaction (ATPM, ATP + H_2_O → ADP + H^+^ + Pi) with *i* constrained to 0 and 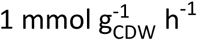, respectively. Maximization of the ATPM reaction, therefore maximization of ATP wasting, entails maximum ATP production (*v*_*ATP*_), since biomass formation is not simultaneously enforced. The objective function values at optimality, i.e., the maximum ATPM fluxes, are used to compute the derivative *d*υ_ATP_⁄*d*υ_i_ representing the ATPsc.

## 3. Results

### 3.1. Coupling chemistries of potential enzyme selection systems span a range of enzyme classes and metabolic sectors

We define an enzyme selection system (ESS) as a microbial chassis strain which allows for the evolutionary optimization of a group of enzymes using ALE experiments and growth as the selection criterion. The reaction catalyzed by each enzyme associated with an ESS is abstracted by one reaction formula, here termed as a coupling chemistry (CC). A CC has an explicit substrate-product pair common to all associated enzymes and a set of generic, enzyme specific reactants (Fig. 1A). For example, a CC with S-adenosyl-L-methionine and its demethylated form S-adenosyl-l-homocysteine as the common substrate-product pair describes many methyltransferases (EC 2.1.1). The generic reactants of the CC thereby represent the methyl acceptor and the methylated product specific to each enzyme. However, CCs do not have to comprise one EC class exclusively but can represent enzymes from multiple different classes if they share the CC’s common substrate-product pair.

**Fig. 1.**
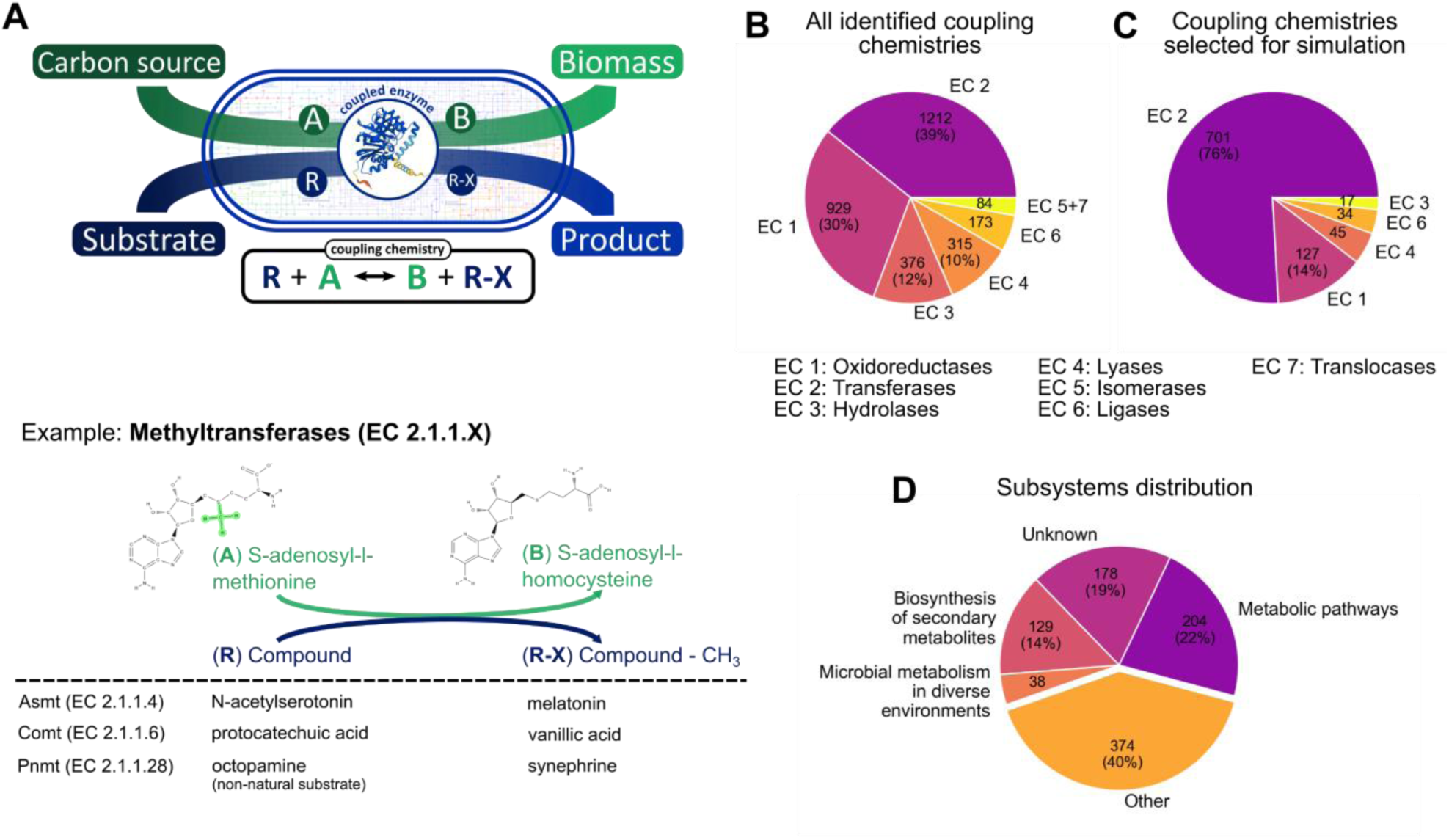
Illustration and analysis of coupling chemistries being generic reaction formulas for groups of enzymes sharing a common substrate-product pair. (A) Scheme of a coupling chemistry (CC) metabolically coupled to growth in a microbial host. All enzymes represented by a CC share the common substrate A and product B which link to the host’s native metabolism. R and R-X are the individual reactants for each enzyme. A methyltransferase reaction scheme is depicted as an example CC for which S-adenosyl-l-methionine and S-adenosyl-l-homocysteine is the common substrate-product pair. The enzymes acetylserotonine O-methyltransferase (Asmt), catechol O-methyltransferase (Comt), and phenylethanolamine N-methyltransferase (Pnmt) and their corresponding reactants are exemplarily shown as members of the methyltransferase CC. Enzyme class distribution among enzymes from all coupling chemistries identified from a BRENDA database search (N=3089) (B) and a subset of those selected for the *in silico* growth-coupling study in *E. coli* (N=924) (C). (D) Distribution of the metabolic sectors associated with enzymes within the coupling chemistries selected for the *in silico* study. A detailed breakdown of the subsystems summarized under “Other” is shown in Fig. S5. Metabolic sector associations were derived from the KEGG database.

To investigate whether ESSs can be designed from generic CCs, CCs comprising two or more enzymes were derived from the BRENDA enzyme database (Jeske et al., 2019). 2564 CCs were identified representing 3089 enzymes spanning the full range of enzyme classes (Fig. 1B). Most CC associated enzymes are oxidoreductases (EC 1) and transferases (EC 2) (together 69%) which reflects the similarity between the CC definition and their catalyzed reactions typically involving a transfer of functional groups, oxygen, or hydrogen atoms from a donor to an acceptor molecule. Methyltransferases (EC 2.1.1) and acyltransferases (EC 2.3.1) were most abundant and represented 10% and 8% of all enzymes assigned to a CC, respectively (Fig. S3).

We selected a subset of all identified CCs for further analysis according to the following criteria: 1) Representing at least five different enzymes to satisfy the demand for a “plug-and-play” evolutionary enzyme optimization platform, and 2) the ability to be integrated into the genome-scale metabolic model iML1515 of *E. coli* K-12 MG1655 (Monk et al., 2017), i.e., their explicit reactants need to be present in the model. As a result, the complete list of CCs was condensed to a core of 44 CCs (Table 2) encompassing 924 enzymes from all top-level enzyme classes but isomerases (EC 5) and translocases (EC 7) (Fig. 1C). As in the complete set of CCs, enzymes covered by the selected CCs primarily belong to oxidoreductases (EC 1) and transferases (EC 2) (together 90%). Despite the bias towards transferases in particular, the selected CCs and their enzymes cover a range of metabolic subsystems according to the enzymes’ pathway associations in the KEGG database (Kanehisa et al., 2017) (Fig. 1D, Fig. S5). As we will show below, the broad coverage of metabolic subsystems can be attributed to the remarkable metabolic versatility and importance of methyltransferases (EC 2.1.1) accounting for a third of all enzymes within the CC subset (Fig. S4).

### 3.2. A workflow for computing enzyme selection system designs for coupling chemistries

An ESS is derived from a CC by growth-coupling the CC’s generic reaction. As is the case for any heterologous reaction, CCs are not necessarily growth-coupled in a wildtype metabolic background. Finding strain designs that enforce a coupling between the catalytic activity of a non-essential or non-catabolic enzyme and biomass formation is a hard task due to the inherent complexity of metabolic networks. Genome-scale constraint-based metabolic models enable a directed, computer-aided search for feasible growth-coupling strain designs and have been widely studied for that purpose. We developed a comprehensive Python-based workflow for finding growth-coupling strain designs, built on the COBRApy metabolic modeling package (Ebrahim et al., 2013) (Fig. 2). At the core of the workflow, the gcOpt optimization algorithm (Alter and Ebert, 2019) provides relevant growth-coupling solutions in the form of appropriate gene knockouts, heterologous reaction insertions, or medium conditions given a metabolic model and reaction database. Briefly, gcOpt solves a Mixed-Integer Linear Programming (MILP) problem that maximizes a minimally guaranteed target reaction flux for a fixed growth rate. Design characteristics such as the growth-coupling strength (GCS) and the maximum growth rate are computed. A detailed description of the workflow is given in the Methods section.

**Fig. 2.**
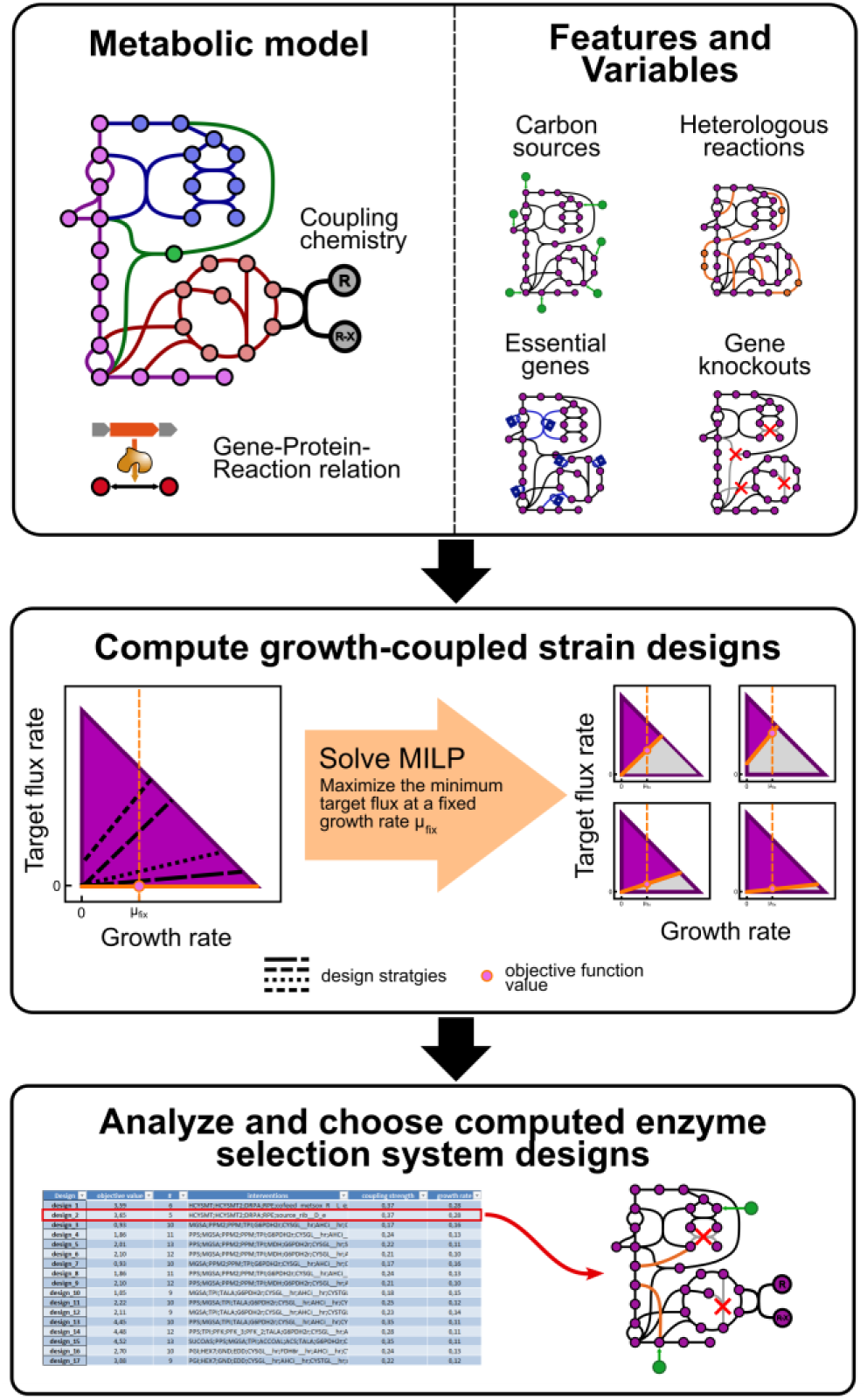
Workflow for computing enzyme selection system designs from coupling chemistries using constraint-based metabolic models. At the first stage, a coupling chemistry is added to a metabolic model, constraints are applied to match the environmental conditions, essential genes are defined, and carbon sources, heterologous reactions, as well as gene knockouts are set as design variables. Secondly, the gcOpt algorithm is applied to derive growth-coupled strain designs, i.e., sets of appropriate design variables, which are being summarized in human-readable tables. At last, suitable strain designs can be chosen for constructing an enzyme selection system considering simulated metadata and experimental expert knowledge.

We applied the growth-coupling strain design workflow to the 44 selected CCs to derive ESSs in *E. coli*. The focus was set on *E. coli* as a platform organism for ESSs, due to the availability of many established genetic engineering tools, its ease of handling in the laboratory, a demonstrated suitability as a multi-purpose microbial cell factory, and its well curated genome-scale metabolic model iML1515 (Monk et al., 2017). To compute a diverse database of potential ESSs, the growth-coupling workflow was applied to each selected CC using multiple optimization problem parameter sets as described in more detail in the Methods section. The parameters that were manipulated included the number of total interventions, heterologous insertions, gene knockouts, as well as the fixed growth rate. Moreover, six variants of the standard iML1515 model and parameter set were employed to investigate the influence of environmental and genotypic conditions on finding growth-coupling strain designs. Details for each applied model variant are summarized in Table 1.

### 3.3. A database of growth-coupling strain designs for enzyme selection system

The ESS design workflow was applied for any combination of selected CC (Table 2) and model variant (Table 1) resulting in a rich database of growth-coupling strain designs (Fig. 3A). Each computed strain design represents a potential strategy for establishing an ESS for the respective CC, and therefore for a specific group of enzymes. 47,635 unique growth-coupling strain designs were identified in total across 33 CCs (77 %) (Fig. 3B, Supplementary Data). For the remaining 11 CCs (33 %) no designs could be identified. The designs show a wide range of GCS values with a median of 0.18, which was used as a cut-off for separating low growth-coupling (GCS < 0.18) from high growth coupling designs (GCS > 0.18) (Fig. 3C). Similarly, maximum growth rates of identified ESS strain designs are widely distributed with a bias towards 0.1 and 0.3 h^-1^ (Fig. 3D). This is a consequence from the fact that the fix growth rate applied in the gcOpt formulation acts like a lower bound on growth for computed strain designs and higher observed growth-coupling strengths generally compromise on maximum growth rates.

**Fig. 3.**
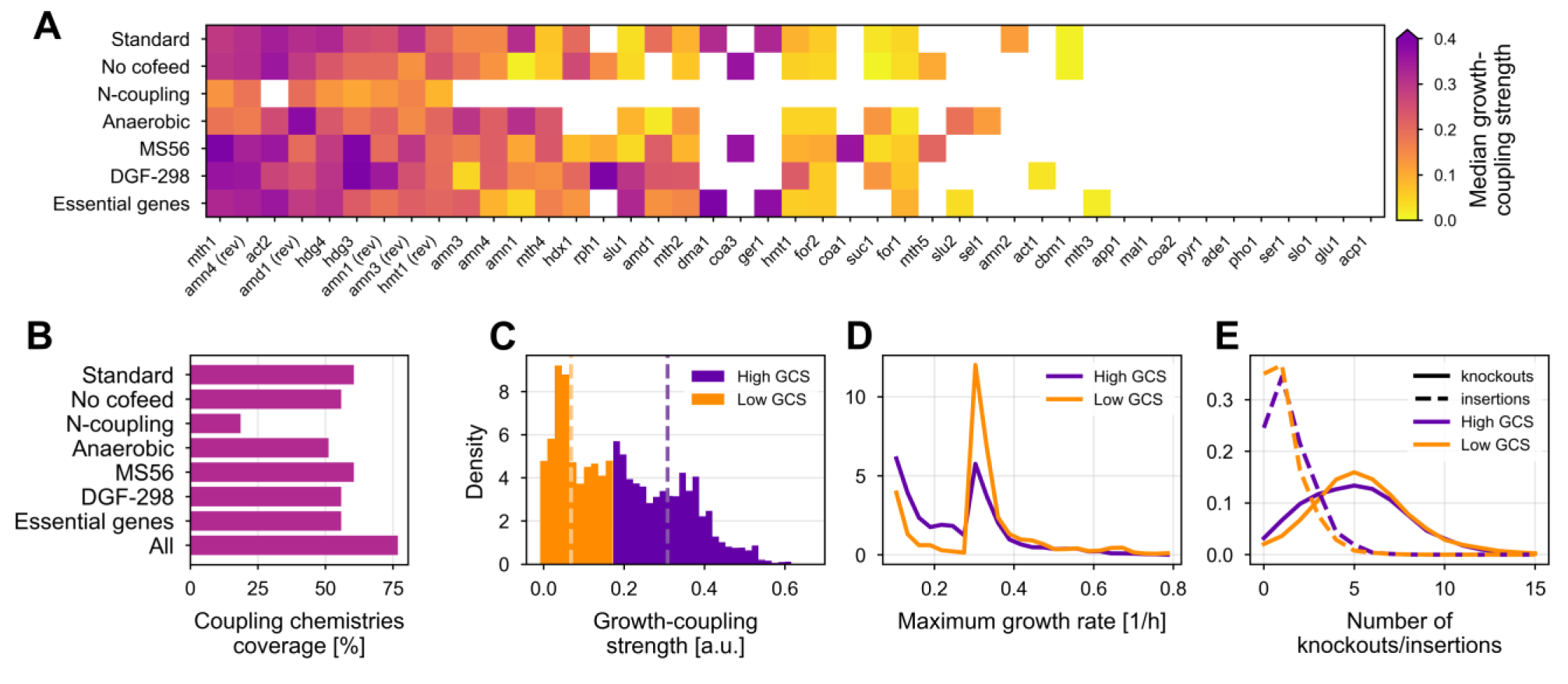
Overview of the strain designs and their basic characteristics encountered for each coupling chemistry (CC). (A) The median growth-coupling strength (GCS) computed from all growth-coupling strain designs for any CC-model combination is shown. White areas denote a complete absence of valid strain design solutions. (B) Share of CCs for which at least one valid growth-coupling strain design was identified using a given model. (C) Distribution density of GCS values for all strain designs identified in this study. The threshold between high (purple) and low (orange) GCS designs was set at the GCS median of 0.18. The dashed lines separately mark the GCS median of all low and high GCS strain designs at 0.07 and 0.31, respectively. Distribution density of the (D) maximum growth rate and (E) the number of knockouts (straight lines) as well as heterologous reaction insertions (dashed lines) for high and low GCS strain designs. Distribution densities were normalized such that the integral for the whole range of each curve equals 1.

### 3.4. Enzyme selection systems are achieved with a reasonable number of interventions

Most computed strain designs and ESSs can be realized with a reasonable number of genetic interventions of five or fewer knockouts (55 %) and two or fewer heterologous insertions (65 %) (Fig. 3E) which is in line with growth-coupling studies of endogenous reactions (Feist et al., 2010; Jensen et al., 2019; Tervo and Reed, 2012). Thus, the implementation of most of the designs in the ESS database is expected to be experimentally feasible. Surprisingly, strain designs with high GCS values require slightly fewer genetic interventions than weakly growth-coupling designs (Fig. S6). 57 % of all high growth-coupling designs, i.e., designs with GCS values greater than the median of 0.18 (cf. Fig. 3C), suggest knockout of five or less genes versus 53 % of designs with low growth-coupling (GCS < 0.18). This counterintuitive discrepancy highlights the importance of the topological position of a target reaction in a metabolic network for finding growth-coupling design strategies and the metabolic burden its activity inflicts on the cellular host.

### 3.5. Genome reduction and medium composition variations only slightly impact the identification of growth-coupling designs

The success rate of finding growth-coupling strain designs for CCs ranges between 51 % and 60 % for most tested model variants and parameter sets (Fig. 3B). Only for the N-coupling case growth-coupling of CCs was largely prohibited, thus rendering the coupling of nitrogen provision from alternative, biotic sources, such as amino acids, to a target reaction as a weak and non-generalizable principle for establishing ESSs. Beside the complete lack of growth-coupling strain designs for 11 CCs, five CCs (rph1, coa3, mth5, coa1, and act1) were only successfully coupled to growth by using genome-reduced models or standard model variants without considering cofeeding of metabolites as design variables. Thus, identification of the strain designs was apparently only made possible by a reduction of the design solution space. The found design solutions for these five CCs nevertheless either show low GCS values, necessitate many interventions, or both, pointing to their principal deficiency for growth-coupling.

Similarly, the use of models representing the genome-reduced *E. coli* strains MS56 (Park et al., 2014) and DGF-298 (Hirokawa et al., 2013) slightly reduces the number of genetic interventions (KS value < 0.12, p-value < 10^-10^, Kolmogorov-Smirnov (KS) test) and knockouts (KS value < 0.17, p-value < 10^-10^, KS test) necessary for growth-coupling CCs (Fig. 4A and B). 20 % of all designs computed with the wildtype *E. coli* model (Standard case) comprise six or fewer interventions compared to 29 % and 30 % of the designs for the MS56 and DGF-298 model, respectively. These differences mainly result from lower numbers of knockouts in the designs for the genome-reduced models. Five or less knockouts are suggested in 49 % of the designs for the wildtype model compared to 62 % and 65 % for the MS56 and DGF-298 model, respectively. On the other hand, the number of heterologous reaction insertions slightly increases (KS value < 0.27, p-value < 10^-10^, KS test) when the genome-reduced models are applied (Fig. 4C). Two or more insertions must be made in 20 % of all wildtype model designs compared to 39 % and 46 % of the MS56 and DGF-298 model designs, respectively.

**Fig. 4.**
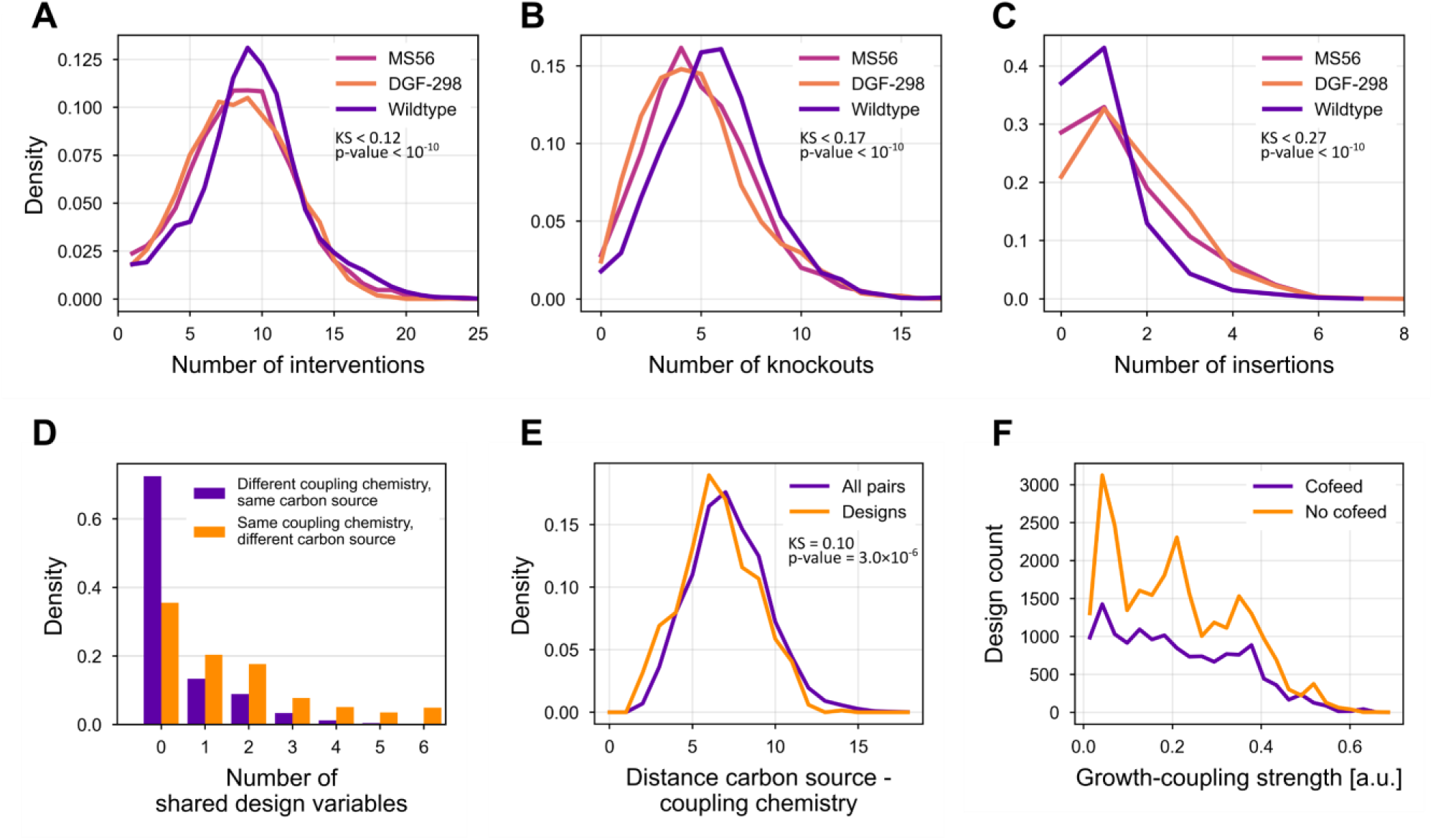
Model- and design variable-specific characteristics of growth-coupling strain designs. Distributions of the number of interventions (A), knockouts (B), and heterologous insertions (C) for strain designs computed using models for the genome-reduced *E. coli* strains MS56 (pink) and DGF-298 (orange), as well as the wildtype strain (purple, Standard model). Upper bounds of KS test statistics and p-values are from Kolmogorov-Smirnov tests between genome-reduced and wildtype strains. (D) The number of shared design variables, i.e., shared knockout and heterologous insertion targets, are shown for 1,000,000 randomly chosen pairs of growth-coupling strain designs. Designs in each pair were computed for different coupling chemistries (CC) but a common carbon source (purple) or for the same CC but different carbon sources (orange). (E) Distribution densities of the distances between the carbon source (extracellular metabolite) and the substrate of a CC. The purple and orange lines show the distributions considering all possible CC-carbon source pairs and only those encountered in computed growth-coupling strain design solutions, respectively. KS test statistic and *p*-value are from a Kolmogorov-Smirnov test. (F) Growth-coupling strength distributions for all strain designs that consider an additional metabolite cofeed (purple) or only a single carbon source (orange). Strain designs for the model variant “No Cofeed” were excluded here.

Independent of the target CC or the chosen model variant, the choice of the primary carbon source does not necessarily restrict the identification of ESS strain designs. Various designs with differing carbon sources were identified for the CCs amenable to growth-coupling. A preference for one or more carbon sources among the designs could not be identified. The choice for a carbon source also does not predetermine the metabolic intervention strategy across CCs. The majority (73 %) of randomly chosen pairs of growth-coupling designs for different CCs sharing the carbon source do not have a single knockout or insertion target in common (Fig. 4D). Similarly, growth-coupling design strategies for the same CC are generally different from each other, if different carbon sources are chosen. 56 % of all screened design pairs of the same CC which do not share the carbon source have either none or one genetic intervention in common. Moreover, the distance in the metabolic network between a carbon source and the substrates of a CC does not appear to be of importance for finding an optimal medium composition for growth-coupling (KS value = 0.1, p-value = 3.0×10^-6^, KS test) (Fig. 4E). The distance distribution for carbon source-CC pairs among the identified growth-coupling designs is comparable to the distribution derived from all possible carbon source-CC pairs. Thus, a possible hypothesis that growth-coupling of a target reaction is more easily established for shorter metabolic distances to the primary carbon source cannot be confirmed based on the design set identified here. Similarly, a switch to anaerobic conditions neither facilitates nor significantly prohibits the identification ESS strain designs across CCs (Fig. 3A and B).

Besides choosing a primary carbon source, addition of a metabolite cofeed was generally allowed as a design variable except for the model variant “No Cofeed”. Examples of previous experimental realizations of growth-coupling systems have frequently applied a cofeed strategy to establish the coupling through an auxotrophy (Harder et al., 2016; Long et al., 2020; Luo et al., 2020, 2019). In principle, if a target reaction is directly involved in the catabolism of the cofed compound, growth-coupling may be established by making essential cellular functions strictly dependent on the metabolite cofeed and therefore the target reaction. This decoupling of essential functions from main metabolic activities promises to generate strong growth-coupling designs while maintaining cellular viability due to the provision of an additional carbon source and the avoidance of rerouting fluxes in central carbon metabolism. However, providing the opportunity for a second carbon source did not fundamentally enable growth-coupling of CCs or increase the GCS of identified strain designs (Fig. 3A, Fig. 4F). If the gcOpt algorithm is allowed to freely add a metabolite cofeed, i.e., designs for the model variant “No Cofeed” are not considered, growth-coupling strain designs without a cofeed predominate. Moreover, the GCS distributions are qualitatively similar for designs with and without a secondary carbon source (Fig. 4F). This observation also applies to the maximum growth rate (Fig. S8). Thus, considering or neglecting a metabolite cofeed does not have a significant effect on the success of finding appropriate growth-coupling strain designs. Addition of a secondary carbon source can still be considered to exhaust all valid strain design possibilities for the final, manual inspection and choice. In summary, a generally valid rule for choosing a carbon source for a given growth-coupling problem cannot be deduced, which highlights the importance of model-based approaches in the decision-making for optimal medium composition and design strategies.

### 3.6. Growth-coupling strength correlates with flux distribution characteristics

As has been argued in the previous sections, the identification of growth-coupling designs for CCs is a challenging task and predictions about the genetic interventions or medium compositions cannot be reliably made *a priori* based on, e.g., the target reaction’s stoichiometry. On the other hand, growth-coupling strain designs follow general metabolic principles, as has been previously shown on designs coupling the secretion of various compounds to growth (Alter and Ebert, 2019). A major finding was that strong growth coupling is enforced by making a target reaction an indispensable part of the major energy (ATP) supply route. The contribution of a target reaction to the cellular ATP supply was quantified by the ATP synthesis capability (ATPsc) measure, which describes the change in ATP production with an enforced unit change in the target reaction flux rate (cf. detailed descriptions in the Methods chapter).

For almost all CC reactions, the ATPsc is negative in the *E. coli* wildtype model (Fig. 5A) indicating that activation of CCs generally exerts a metabolic burden on the cells by diverting fluxes away from optimal ATP and biomass synthesis routes. Applying the identified ESS strain designs increases the ATPsc to positive values across CCs (KS value = 0.74, *p*-value < 10^-10^) (Fig. 5A, Supplementary Data) entailing that the metabolic network is reconfigured such that the activity of CCs positively contributes to the ATP supply. Furthermore, positive linear correlations between ATPsc and GCS are observed for a majority of CCs (91 %) of which 80 % are significant (*p*-value ≤ 0.01, Pearson correlation) according to the Pearson correlation coefficient (Fig. 5B, Fig. S9). Thus, there is a quantitative link between the coupling strength and the contribution of a CC to cellular ATP supply.

**Fig. 5.**
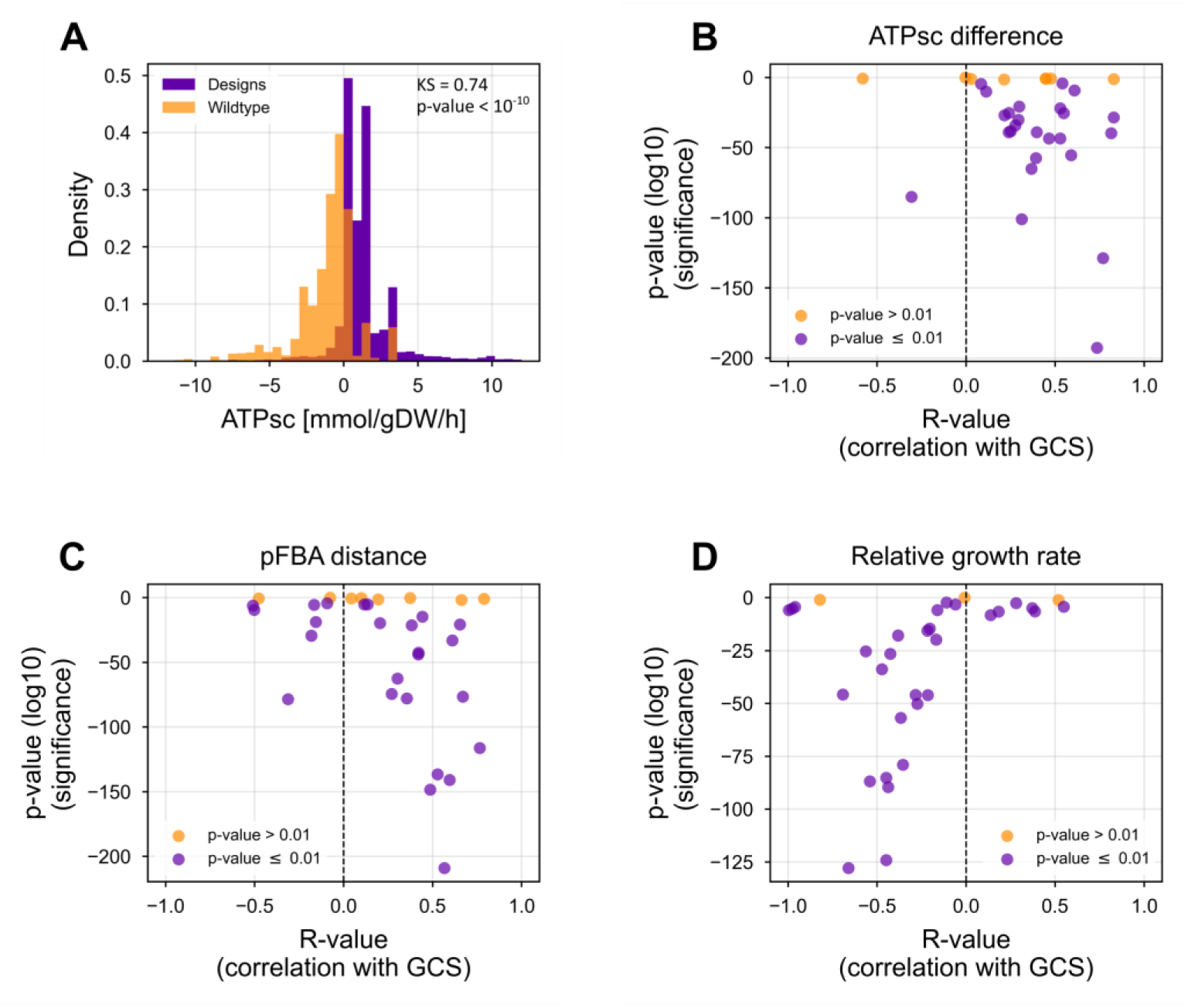
(A) Absolute ATP synthesis capability (ATPsc) distributions for the generic stoichiometric reactions of all coupling chemistries (CC) in a wildtype metabolic model (orange) and in models with applied growth-coupling strain designs identified in this study (purple). KS test statistic and *p*-value are from a Kolmogorov-Smirnov test between the wildtype and design distribution. Several quantitative metabolic measures were computed for all identified strain design for a CC and linearly correlated with the growth-coupling strength (GCS) (B-D). The goodness-of-fit, i.e., the Pearson correlation coefficient (R-value), for each individual CC and the corresponding significance (*p*-value) are shown. The quantitative metabolic measures are (B) the difference of ATPsc values for a CC, (C) the distance between flux distribution vectors of pFBA solutions at optimal growth, and (D) the change in maximum growth rates between a wildtype and strain design model for a given design and CC.

Slightly less prominent observations were made for the distances between flux vectors of parsimonious Flux Balance Analysis solutions (pFBA) (Lewis et al., 2010) and corresponding growth rates computed for wildtype and growth-coupling strain design models at optimal growth (Fig. 5C and D, Fig. S10 and Fig**. S11,** Supplementary Data). For 55 % of the CCs, the distance of pFBA solutions between growth-coupling designs and wildtype significantly increases with higher coupling strengths (*p*-value ≤ 0.01). Thus, for approximately half of the CCs, the implementation of ESS designs with increasing coupling strengths is associated with an increased need for significant metabolic flux rearrangements. Even though for the other half the extent of flux rearrangements does not appear to be related to the choice of the strain design, for the majority (73 %) of CCs, tighter coupling in terms of higher GCS values significantly leads to increasingly severe growth defects.

### 3.7. Identified enzyme selection systems cover a range of pathways relevant for bioproduction purposes

Although a robust ESS could not be identified for every CC, we still expect that a wide range of molecules can be growth-coupled using the designs proposed here. Especially since some CCs represent a multitude of unique, known reactions, there is potential for the development of biocatalysts for various distinct, industrially relevant products. To further explore this notion, we set out to map which compounds could be produced through a pathway of at most eight heterologous enzymes that ends with a viable CC, starting from *E. coli* native metabolites. The choice of native metabolites as a starting point may exclude reactions that modify complex, non-native compounds into industrially relevant products, but it ensures a focus on whole-cell biocatalysis, where the product of interest can be synthesized from any of the carbon sources the cell is able to utilize.

First, reactions matching the templates of viable CCs (Table 2) were identified in the KEGG database of biochemical reactions (Kanehisa et al., 2017). Subsequently, the KEGG database was traversed in a retrosynthetic manner, using a maximum of eight steps, to arrive at metabolites endogenous to *E. coli* (Zhang et al., 2016), as determined by their presence in the iML1515 metabolic model (Monk et al., 2017). This methodology readily revealed 285 products that have the potential to be produced using an ESS. Since there was no lower bound imposed on the number of heterologous reactions, these products include 119 compounds already present in the *E. coli* metabolome. Generally, a large fraction of the amino acids and organic acids in the central carbon metabolism is used as a starting point for the heterologous biosynthetic pathways, in addition to the cofactors and more trivial compounds, such as H^+^, O_2_ and H_2_O.

Most products are associated with transaminase and methyltransferase reactions (Fig. 6A), where the mth1 (S-adenosyl-L-methionine-dependent methyltransferase) CC accounts for 93 products. For 84% of the products, a single CC was identified, with the remaining compounds largely representing endogenous metabolites involved in a vast number of reactions. Among the products associated with just a single CC, a large fraction (95%) is non-native to *E. coli*. The redundancy of CCs for non-native compounds all occurred within the same class of chemistries (mainly transaminases), whereas compounds endogenous to *E. coli* sometimes span multiple classes.

**Fig. 6.**
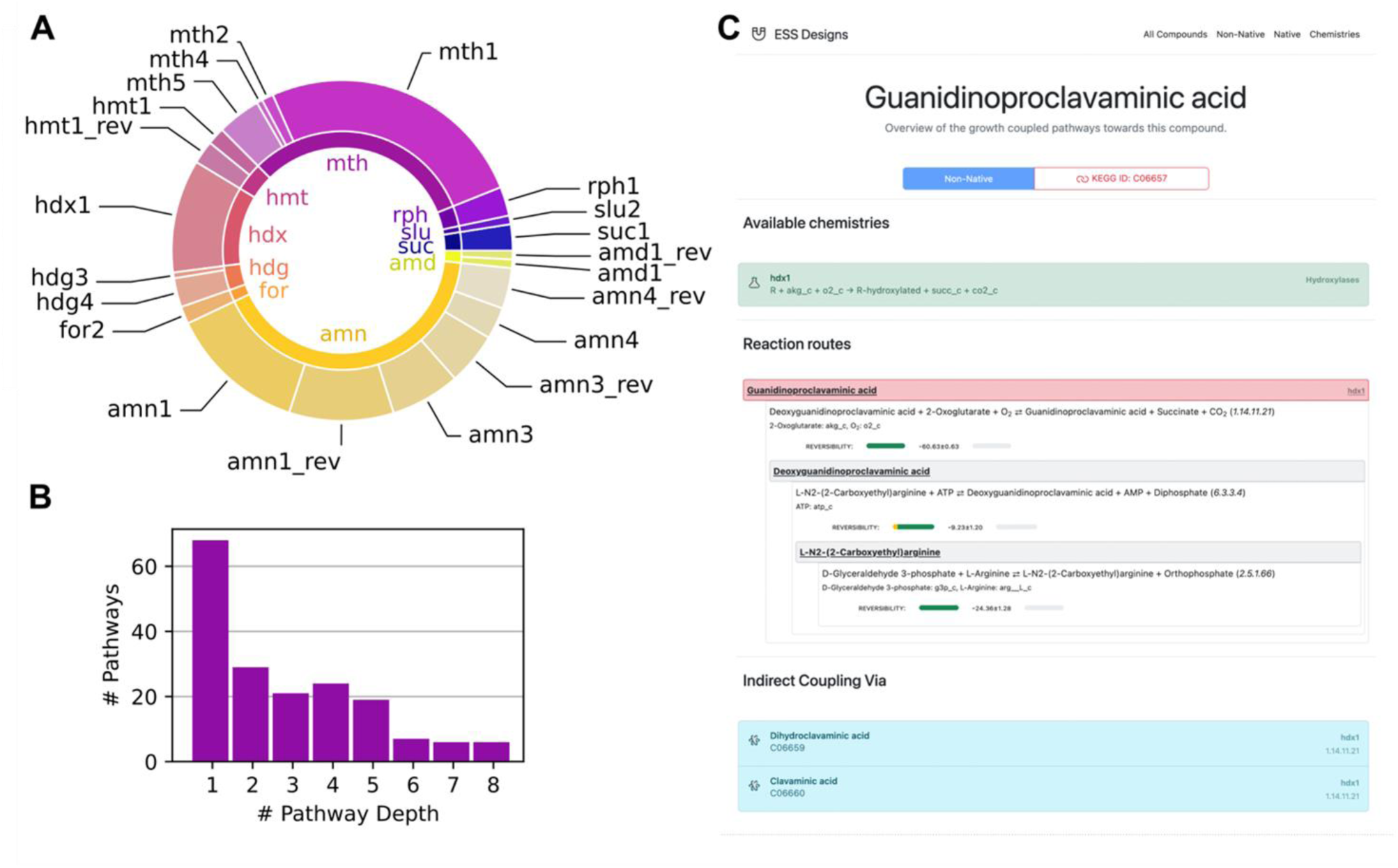
(A) Classification of the pathways identified in a retrosynthetic analysis according to associated coupling chemistries (CCs). The inner ring of the donut chart represents the classification of the pathways into the various enzyme classes, whereas the outer ring specifies their subdivision into CC IDs (Table 2). (B) Distribution of pathway lengths, as measured by the maximum number of heterogenous reaction steps between a native *E. coli* metabolite and the non-native pathway product. The search depth of the retrosynthetic approach was limited to eight reaction steps. (C) Screenshot of a product information page of the ESS designs database (https://biosustain.github.io/ESS-Designs/). The page displays relevant CC information, the heterogenous pathway towards the product, and indirect coupling approaches, when applicable.

We determined the depth of each pathway leading to a non-native product in *E. coli* and observed a notable trend wherein the number of pathways decreased as the reaction depth increased (Fig. 6B). Notably, loops were disallowed in the retrosynthetic analysis, preventing the overrepresentation of longer pathways due to, e.g., trivial back-and-forth traversals over reversible reactions. Given that almost two-thirds of the non-native products were reached only after at least two consecutive heterologous reaction steps, there is a wide array of downstream intermediates whose production can be optimized indirectly through selective pressure on the growth-coupled upstream product. Inclusion of this set of compounds adds 147 novel products to our analysis and provides an alternative, indirect coupling option for 45 already included products. Implementation and analysis of data obtained for indirectly growth-coupled products requires careful consideration, since not all adaptations necessarily benefit the production of the target compound and transport as well as toleration of the compound are not directly part of the growth-coupling mechanism.

An easily navigable instance of computed ESS strain designs as well as associated pathways and products is available at https://biosustain.github.io/ESS-Designs/ (Fig. 6C). Access is provided to a subset of ESS strain designs computed in this work due to license terms. These include 25,505 unique designs for major enzyme classes such as methyltransferases and transaminases, among others. The online database can be searched by CCs or target compounds yielding suggestions for ESS strain designs computed in this study and growth-couplable pathways to the queried product, respectively. We further enriched the dataset with reaction reversibility estimates, as determined using the eQuilibrator web interface (Beber et al., 2021), and design features, such as the predicted maximum growth rate, to facilitate a direct assessment of the suitability of strain designs for a given application. Thus, a rich and accessible resource is provided for engineering cellular enzyme optimization platforms with an application focus for industrial biotechnology purposes.

## 4. Discussion

The development of microbial cell factories and cost-effective bioprocesses hinges on the availability of heterologous enzymes with high catalytic efficiencies towards the production of a desired compound. Here, we introduce the concept of enzyme selection systems (ESSs) as a whole-cell tool for improving enzyme properties and screening of environmental conditions on enzyme activity by using growth as a selection criterion. Besides the computational tools and workflows for individually deriving ESS engineering strategies, a database of designs for multi-enzyme engineering platforms is provided. These platforms give access to pathway engineering capabilities for the synthesis of various biotechnologically relevant compounds without the need for specialized laboratory equipment.

The first proof-of-principles for ESSs were conducted by Luo et al. (2020, 2019) and basic parts of the experimentally implemented strain designs were identified by the computational approach presented in this study. In the case of the ESS for methyltransferases (mth1, Table 2), model-based predictions suggest, among other strategies, the same medium composition (glucose and methionine as carbon sources) and heterologous insertions (cystathionine-β-synthase, cystathionine-γ-lyase), but differing gene knockouts as compared to the manually derived design. This is due to the fact that the experimentally knocked out gene *cysE* (b3607) is deemed essential according to the essentiality dataset and was therefore excluded as a design variable. However, even by disregarding gene essentialities, the growth-coupling strategy from Luo et al. (2019) was not exactly reproduced, which is explained by the particularly low growth-coupling strength (GCS) of their design (0.004, Fig. S12). Due to this, the growth-coupling design algorithm skipped it in favor of strategies with higher GCS values. Given the successful application of the ESS for producing improved methyltransferase variants, weak coupling designs should not be generally dismissed as discussed in more detail below. Consequently, it is necessary to test diverse sets of model parameters and constraints to exhaustively compute growth-coupling designs. This provides the user with full flexibility in selecting an appropriate strategy. Nevertheless, rigid model parameterization, e.g., by using conditional experimental data, can ensure experimental feasibility while facilitating the computation of diverse growth-coupling strategies, including simple and tractable designs. To generally encourage and guide the implementation of ESSs, we discuss the implications for design principles and the implementation process based on the properties of identified designs in the following.

### 4.1. Enzyme selection system designs show shared metabolic principles, but their identification eludes simple rules

The observed diversity of growth-coupling strain designs for coupling chemistries (CCs) yielding ESSs suggests that deriving growth-coupled systems or increasing the coupling strength is not necessarily restricted to complex genetic engineering strategies involving a multitude of interventions such as gene knockouts or heterologous reaction insertions. Ultimately, the involvement of the reactants of an enzymatic target reaction within the microbial host’s metabolic network determine if a target enzyme’s activity can be effectively coupled to growth. For example, the success rate of finding growth-coupling designs and the coupling strength for CCs transferring low-molecular groups, e.g., methyltransferases (mth1-5), transaminases (amn1-4), or dehydrogenases (hdg3), was relatively high compared to CCs such as CoA-transferases (coa1-3), glutamate ligases (glu1), or malonyl transferases (mal1). In fact, among the 11 CCs (23 %) for which no valid growth-coupling designs were found, CoA, malonyl, phosphate, or 3-amino-3-carboxypropyl groups are involved among others in the reaction stoichiometries highlighting the difficulty to growth-couple reactions transferring or otherwise acting on large or highly connected molecular groups and cofactors. Even if designs were found for CCs involved in the transfer of high-molecular groups, e.g., carbamoyl (cbm1) or pyruvate (pyr1), these generally require an increased number of interventions (Fig. S7) or show low GCS values.

However, a strict rule for predicting the success of growth-coupling a CC or the design itself based on the reaction stoichiometry could not be deduced. For example, neither the connectivity of reactants nor their molecular weights were found to correlate with the ability to growth-couple a corresponding CC or the growth-coupling strength of identified strain designs. Thus, designing ESSs or other growth-coupled systems eludes intuition and requires a model-based optimization approach. On the other hand, we have demonstrated that the establishment of ESSs for CCs, if amenable to growth-coupling, is flexible in the choice of medium composition, including oxygen availability. This flexibility allows ESS design strategies to be tailored to a given biotechnological application and its process requirements.

For the CCs amenable to practical growth-coupling strategies, the complexity of strain designs was further decreased in genome-reduced strains. Thus, the implementation of ESSs can benefit from genome reduction in terms of engineering efficiency. This potential may be enhanced by creating genome reduced strains with a minimal metabolic repertoire. So far, metabolic network-agnostic approaches were used for deducing genome reduction strategies by focusing on the deletion of insertion sequences or large clusters of non-essential parts of the genome, as was the case for MS56 (Park et al., 2014) and DGF-298 (Hirokawa et al., 2013). A strategy that precisely and specifically reduces non-essential and redundant parts of the metabolic network while accounting for synthetic lethality promises to advance genome reduction in *E. coli* and reduce the number of interventions necessary to establish growth-coupling designs.

While the process of deriving ESS strain designs is non-trivial, the intertwining of the activity of a target reaction with cellular energy metabolism was identified as a general principle governing ESS strain designs. Although an interdependence between ATP supply and target reaction activity may seem trivial for a growth-coupled system, it has fundamental implications for the metabolic principle of growth-coupling strain designs: Since ATP is the central energy currency and is involved in many cellular processes, growth-coupling of CCs is imposed by introducing a strict link between the target reaction and metabolic activity as a whole rather than the synthesis of just one or a few biomass precursors. This is in line with growth-coupling principles derived from designs coupling various metabolite secretion pathways to growth (Alter and Ebert, 2019).

### 4.2. Implementation of ESSs requires a trade-off between growth-coupling strength and growth rate

The analysis of identified enzyme selection system (ESS) strain designs in this work has implications for the implementation of ESSs and growth-coupling strategies in general. It has been recognized that high coupling strengths are desirable in growth-coupling systems to guarantee an evolutionarily robust activity of a target functionality (Legon et al., 2022), i.e., enzymatic catalytic activity in the case of ESSs. High growth-coupling strengths (GCS) imply a strong selection pressure on the growth-coupled target enzyme’s activity since it must carry high catalytic flux per growth rate unit. This enables better control of the selection pressure through the target enzyme expression strength, while avoiding an increased metabolic burden on the ESS host imposed by an over-allocation of cellular resources toward enzyme expression. Moreover, a tighter growth-coupling results in a more direct, and therefore more accurate, readout of the target enzyme’s activity based on the growth rate of the ESS, which is useful in applications where the ESS is used as an enzyme activity screening system.

However, higher coupling strengths divert (carbon) flux distributions away from optimal energy, redox, and biomass precursor generation and towards the target reaction, thus compromises cellular viability. Furthermore, model-based phenotype predictions are usually only reproducible in evolved strains (Fong and Palsson, 2004; Ibarra et al., 2002). Thus, the experimental cellular viability of ESSs is expected to be even lower than the *in silico* predictions, as was demonstrated by a metabolic model-based investigation of a genotype-phenotype dataset of growth-coupling strain designs extracted from literature (King et al., 2016). Immoderate growth defects prior to laboratory evolution can hamper applicability of ESSs and growth-coupling designs in general, particularly if a high initial growth rate is required for applying an ESS in an ALE experiment. Besides ALE, artificially generated enzyme mutant libraries can also be applied to ESSs, thus circumventing the need for a minimum cellular viability and allowing for augmenting the selection process with knowledge from previous structural analyses or specifically increasing the mutation frequency.

In summary, ESS strain designs enabling only suboptimal growth-coupling still have their justification for practical implementations. The diversity of computed growth-coupling strain designs in this work allows for matching the design strategy to the desired strain properties, e.g., to enable a reasonable coupling strength between the target enzyme and growth while maintaining high cellular viability.

### 4.3. Enzyme selection system designs can escape growth-coupling through unknown enzymatic activities

Although model microorganisms, such as *E. coli,* have been intensively studied for decades, our knowledge about their enzymatic and metabolic repertoire is generally incomplete or not fully covered by metabolic models. For an implemented ESS design, the upregulation of unknown, shallow, or otherwise suppressed cellular functions can cause a metabolic escape, allowing the cell to bypass the coupling and evolve to higher growth rates without improving the target enzyme activity. Such functions can include enzyme promiscuity toward an alternative substrate or the metabolic activity of an unknown enzyme. A previous study has shown that even low-level promiscuous enzyme activities are able to sustain novel phenotypes if evolved and accentuated in ALE experiments (Guzmán et al., 2019). Thus, a high selective pressure on engineered strains is likely to evoke metabolic escapes from growth-coupling of the CC and the corresponding enzyme during ALE experiments. An emerged metabolic escape can be mitigated by eliminating the upregulated, unwanted function or gene. For strain designs with a strong growth-coupling strength and thus a high chance of triggering a metabolic escape from growth-coupling, repeated design-build-test-learn (DBTL) cycles are likely required for establishing a robust ESS for a given CC or enzyme. While the process of mitigating metabolic escapes delays the construction of an ESS, it offers opportunities for metabolic discoveries and the improvement of metabolic models as a byproduct. The emergence of metabolic escapes triggered by unknown cellular functionalities may be avoided in the first place by using genome-reduced strains as ESS hosts. By applying a host strain lacking non-essential genomic parts, the chance for metabolic escapes caused by an evolved upregulation of uncharacterized metabolic functions is reduced. Consequently, the need for feedback loops between design and testing of ESS strains is reduced.

## 5. Conclusion

A general concept was introduced for the construction of microbial chassis strains, so called enzyme selection systems (ESS), which allow for a directed *in vivo* engineering of a class of enzymes using adaptive laboratory evolution (ALE) experiments. The concept is based on coupling chemistries (CC), i.e., generic reactions representing multiple enzymes of a particular class. A large-scale *in silico* study was conducted to identify growth-coupling strain designs for selected CCs and to build a database of ESSs for a broad range of enzymes. Although ESS designs could not be determined for all selected enzymes classes or CCs, the ESS database still covers a wide range of metabolic pathways and associated compounds, thus opening the possibility for in-place pathway engineering approaches for many applications. It was shown that practical, experimentally accessible growth-coupling designs generally exist, and no specific requirements are placed on the microbial base strain or the experimental conditions. Thus, there is a certain degree of freedom in choosing a favorable carbon source and metabolic engineering strategy and this study provides the database for engineering experts to make suitable choices. Moreover, the provided computational tools and workflow enable the comprehensive search for growth- or flux-coupling design strategies for any given application or microbial host. The decision-making process for a growth-coupling design will always involve a compromise between viability of the mutant strain or ESS and the growth-coupling strength. Future experimental efforts will reveal the influence of the coupling strength, i.e., the level of selection pressure on a target gene, on the evolution of improved target gene variants and its optimal balance with inevitable growth defects induced by design. For example, high ESS viability and prevention of metabolic escapes from growth-coupling, both assured by low coupling strength design choices, may be more advantageous for productive ESSs than strongly coupled systems with heavily rerouted metabolic fluxes, which are likely to develop unforeseeable phenotypes. Considering the power of the first experimental examples of ESSs (Luo et al., 2020, 2019), the presented concept of ESS, the corresponding design database, and the computed characteristic features of the designs within, are a cornerstone for establishing ESSs as a tool for enzyme engineering purposes.

## Acknowledgments

The authors acknowledge funding from the Novo Nordisk Foundation (NNF20CC0035580) and thank Lei Yang and Jinbei Li for fruitful discussions of this work from an experimental point of view.

## Contributions

Conceptualization: Tobias B. Alter, Colton J. Lloyd, Adam M. Feist, Bernhard O. Palsson, Daniel C. Zielinski

Investigation and Visualization: Tobias B. Alter

Formal analysis: Tobias B. Alter, Pascal A. Pieters

Software: Tobias B. Alter, Colton J. Lloyd, Pascal A. Pieters

Supervision: Emre Özdemir, Bernhard O. Palsson, Daniel C. Zielinski

Writing – original draft: Tobias B. Alter, Pascal A. Pieters

Writing – review & editing: all authors

## Abbreviations

LE: adaptive laboratory evolution
ATPsc: ATP synthesis capability
CC: coupling chemistry
ESS: enzyme selection system
FVA: flux variability analysis
GCS: growth-coupling strength
GPR: gene-protein-reaction
KS: Kolmogorov-Smirnov
MILP: mixed integer linear program
pFBA: parsimonious flux balance analysis

