## Supplemental Figures and Tables for "Metabolic growth coupling strategies for *in vivo* enzyme selection systems"

**Supplementary Material**

**Supplementary Data.** The Supplementary Data Excel file contains detail information and data used for or generated by the computational approaches presented in this manuscript.

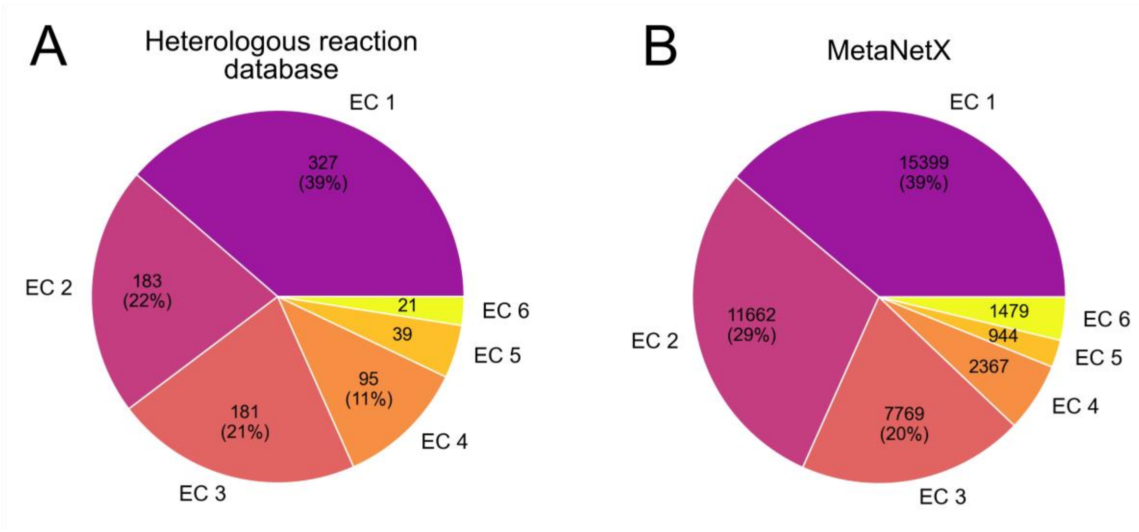


**Fig. S1.** EC number distributions of reactions and corresponding enzymes contained in (A) the heterologous reaction database for the metabolic model iML1515 and (B) the MetaNetX database (version 4.4). The total number of reactions in the heterologous reaction database is 1298 of which 846 (65%) were associated to an EC number. The total reaction count on MetaNetX is 39620. Translocases (EC 7) have been excluded for visualization purposes (N=115, 0.3%).

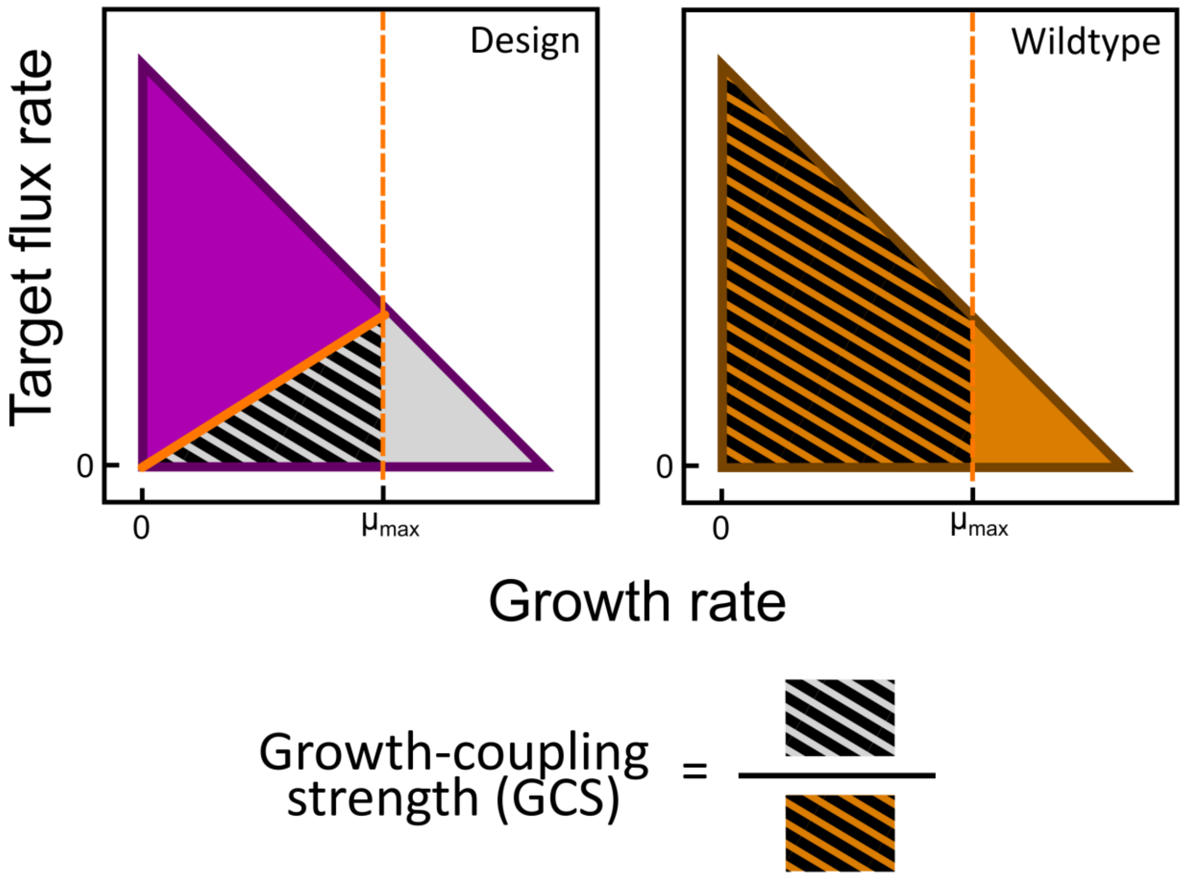


**Fig. S2.** Schematic calculation of the growth-coupling strength (GCS) based on 2D projections of the target flux space span by constraint-based metabolic models. The purple and orange area are the flux spaces accessible by the model with an applied growth-coupling strain design and the wildtype, respectively. The gray area is the flux space accessible to the wildtype model but being made inaccessible by the growth-coupling design. The GCS is computed by dividing the inaccessible flux space area below the lower flux boundary in the in the design model (hatched gray) and the accessible area in the wildtype model up to the maximum growth rate $\mu_{max}$ of the design model (hatched orange).

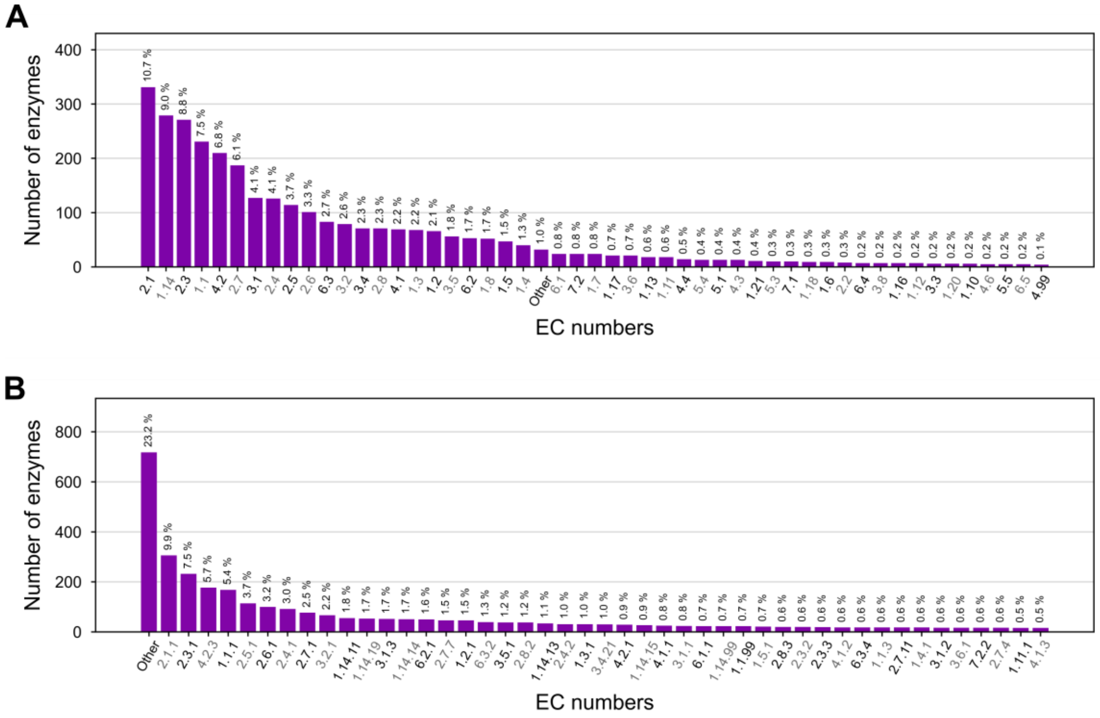


**Fig. S3.** Distribution of EC numbers on the second (A) and third (B) digit level among enzymes associated with one of all 2564 identified coupling chemistries. All EC numbers representing less than 0.1 % (A) or 0.5 % (B) of the total set of CC-associated enzymes are summarized in “Other”, respectively.

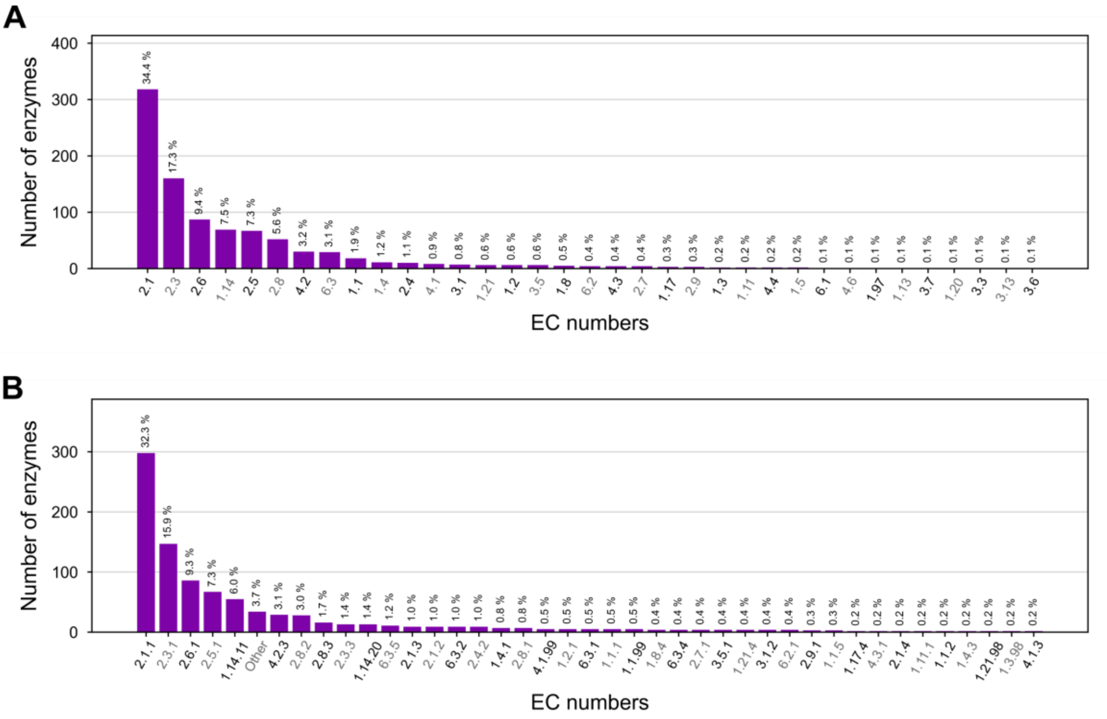


**Fig. S4.** Distribution of EC numbers on the second (A) and third (B) digit level among enzymes associated with one of the 44 selected coupling chemistries (CC). In subfigure (B), all EC numbers representing less than 0.2 % of the selected set of CC-associated enzymes are summarized in "Other".

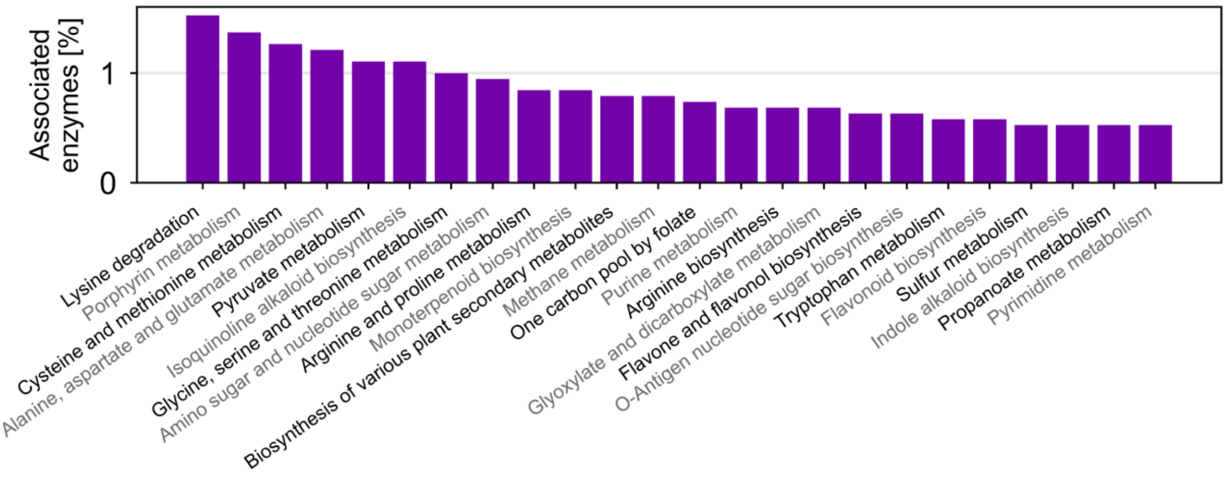


**Fig. S5.** Extract of the distribution of the metabolic sectors associated with enzymes within the 44 coupling chemistries selected for the *in silico* study. Only those metabolic sectors were considered which have associations with less than 4 % of the considered enzymes.

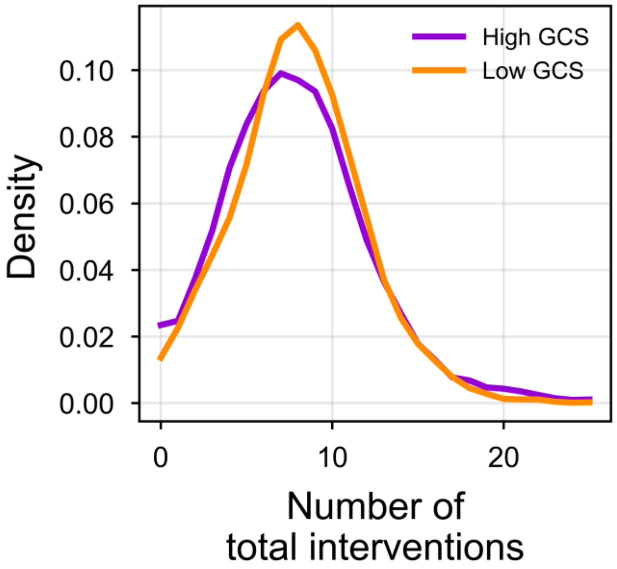


**Fig. S6.** Distribution of the total number of interventions including knockouts and heterologous insertions among all identified growth-coupling strain designs showing a high (purple, > 0.18) or low (orange, < 0.18) growth-coupling strength (GCS), respectively.

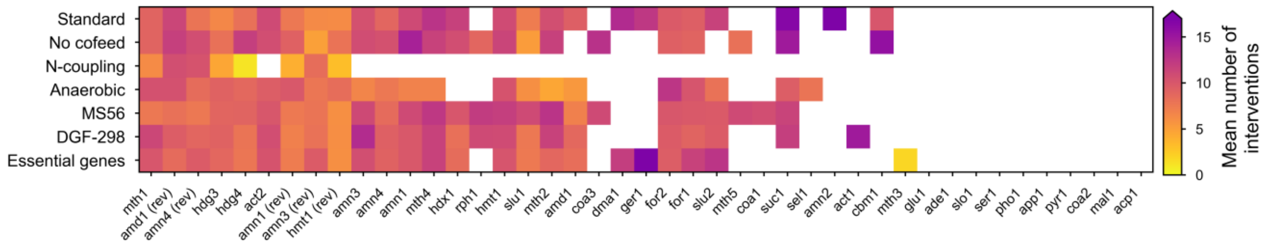


**Fig. S7.** The mean number of interventions including knockouts and heterologous insertions of all growth-coupling strain designs for any coupling chemistry-model combination is shown. White areas denote a complete absence of valid strain design solutions.

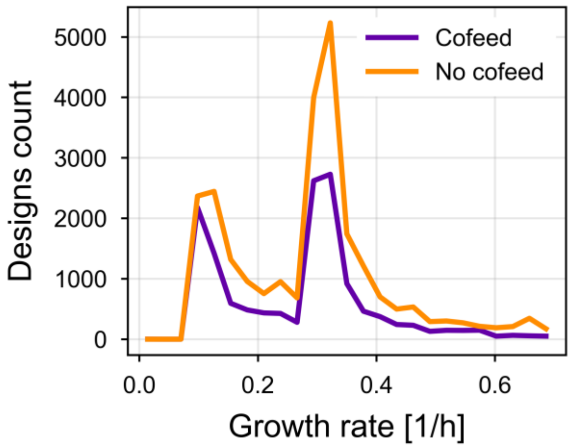


**Fig. S8.** Maximum growth rate distributions for all identified growth-coupling strain designs that consider an additional metabolite cofeed (purple) or only a single carbon source (orange). Strain designs for the model variant “No Cofeed” were excluded here.

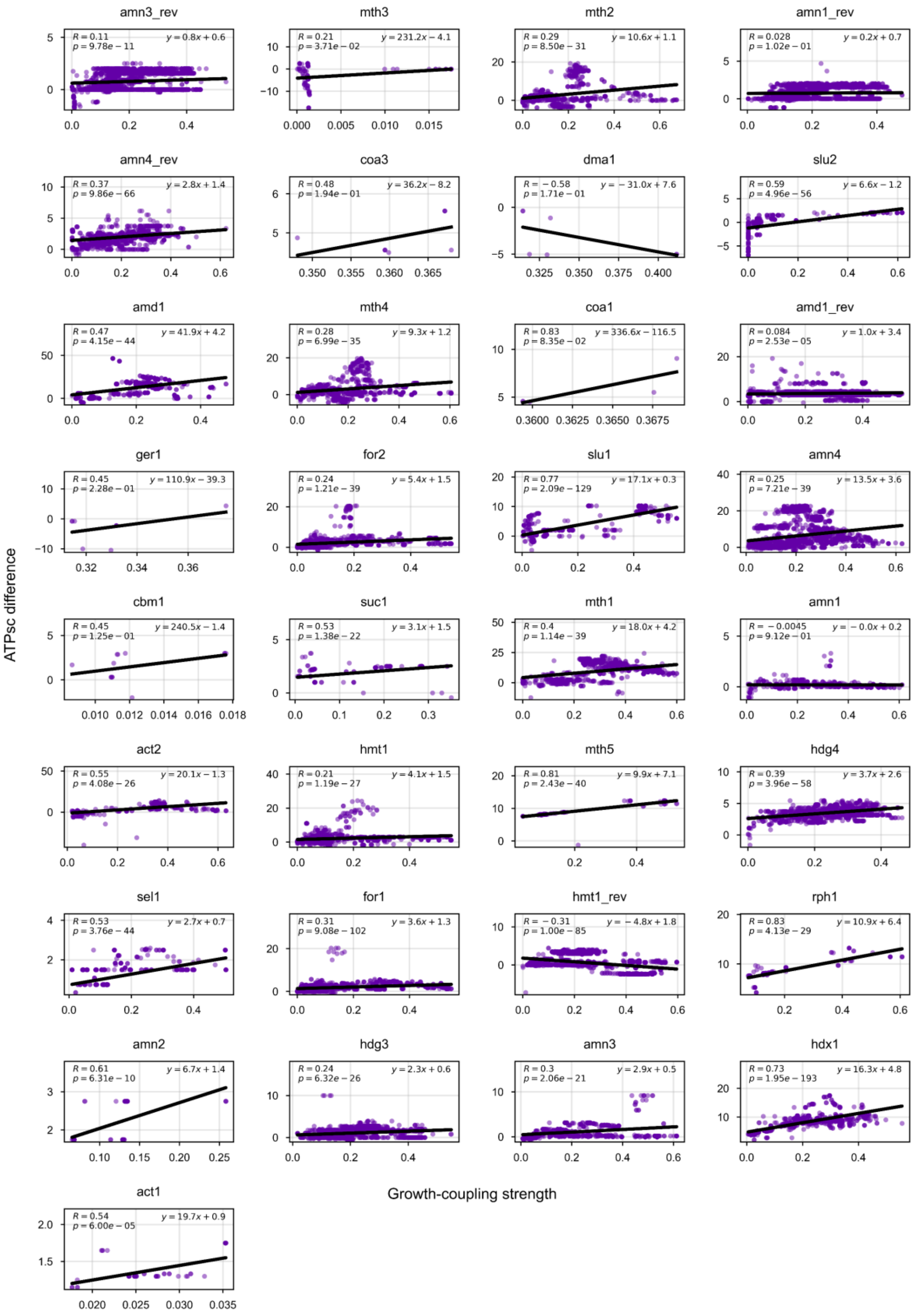


**Fig. S9.** Correlations between the growth-coupling strength and the ATPsc difference of identified growth-coupling strain designs for each coupling chemistry. ATPsc difference denotes the change in ATPsc values between the wildtype *E. coli* model and in the model with an applied growth-coupling design strategy. Linear regressions, corresponding fitted linear equations, correlation coefficient (R-value), and the significance of the correlations (*p*-value) are shown.

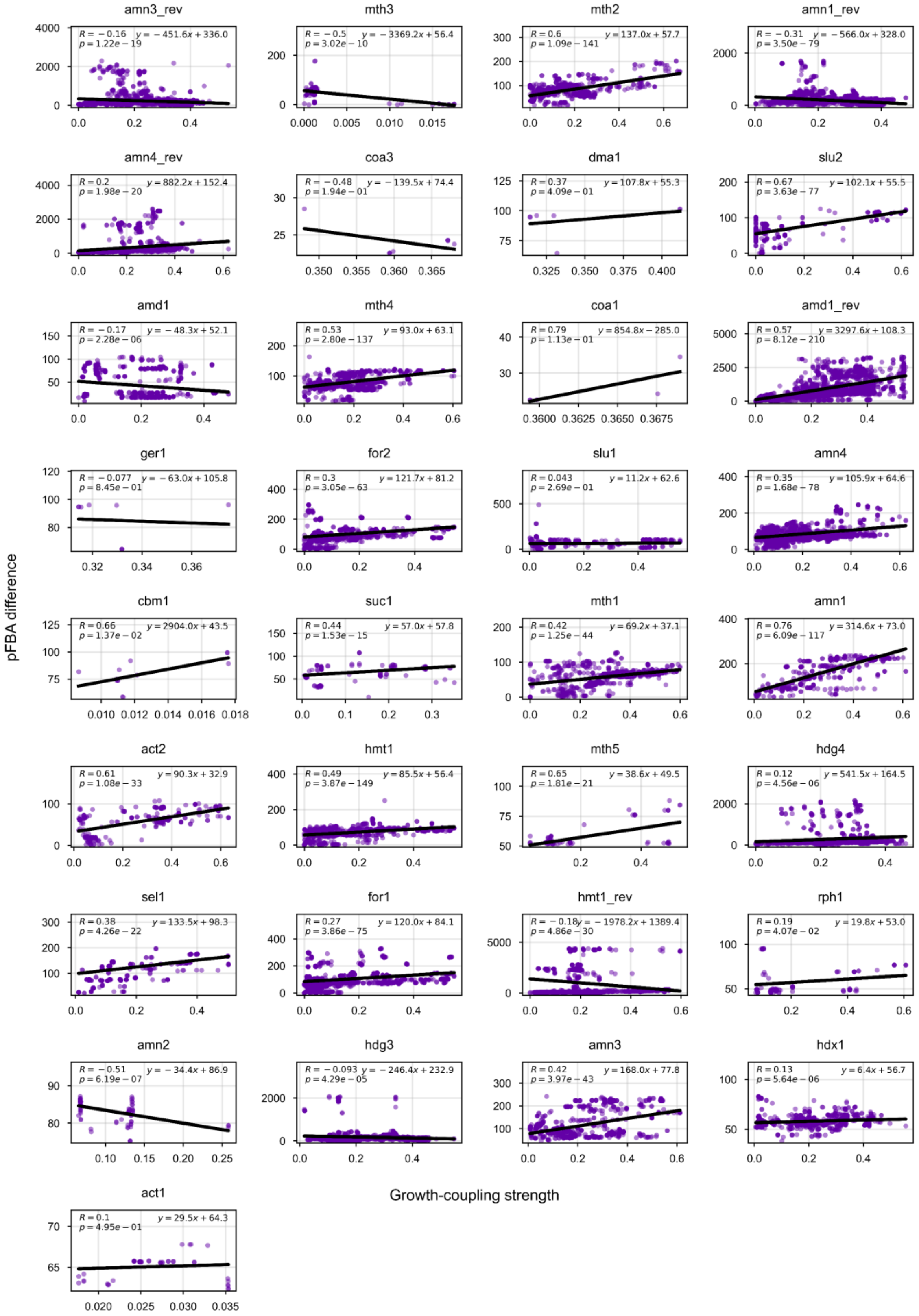


**Fig. S10.** Correlations between the growth-coupling strength (GCS) and the parsimonious Flux Balance Analysis (pFBA) solution distance between wildtype and identified growth-coupling strain designs for each coupling chemistry. Linear regressions, corresponding fitted linear equations, correlation coefficients (R-value), and the significance of the correlations (*p*-value) are shown.

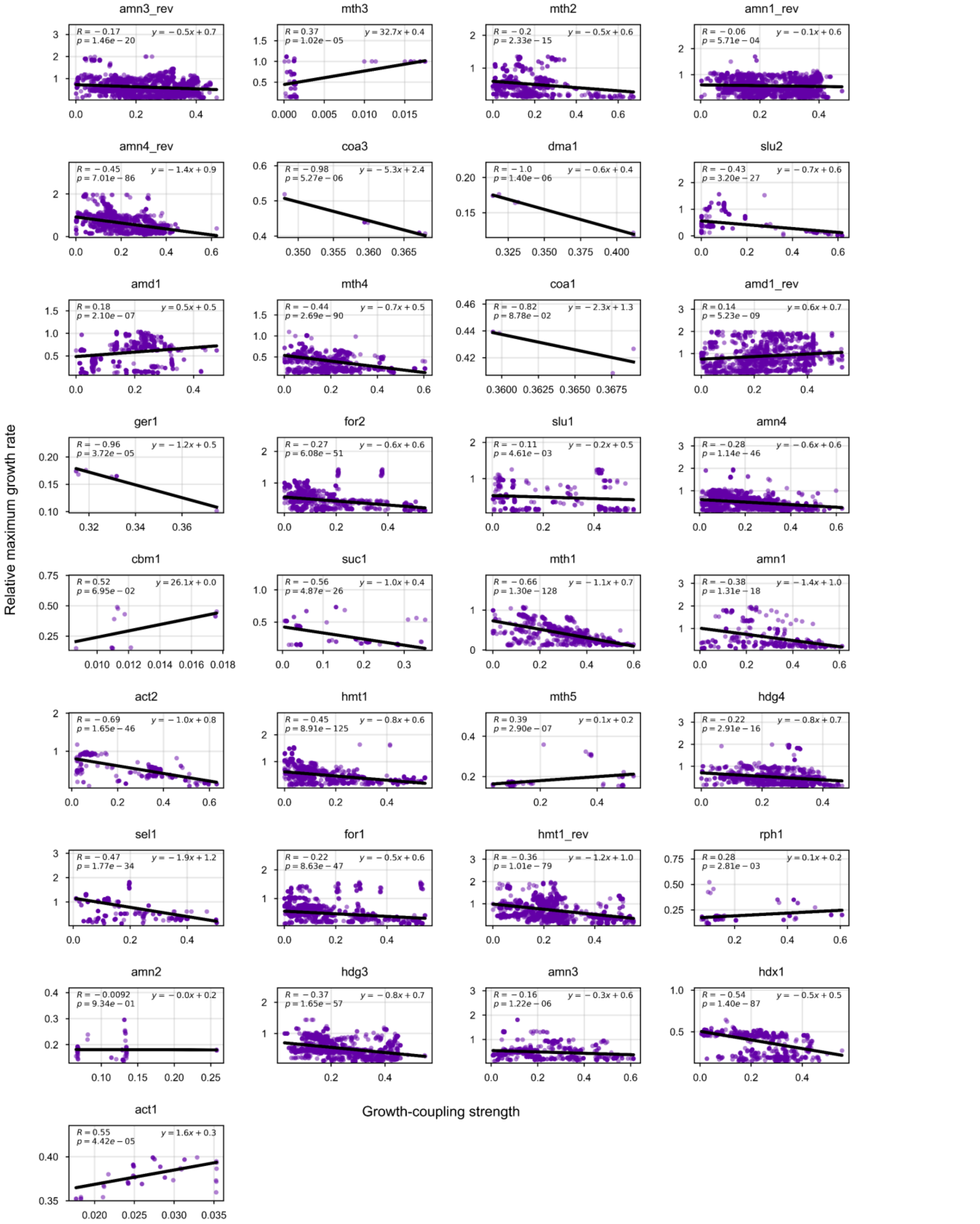


**Fig. S11.** Correlations between the growth-coupling strength (GCS) and the maximum growth rate for growth-coupling strain designs relative to the maximum growth rate of the wildtype model for each coupling chemistry. Linear regressions, corresponding fitted linear equations, correlation coefficients (R-value), and the significance of the correlations (*p*-value) are shown.
